# A neuroepithelial wave of BMP signalling drives anteroposterior specification of the tuberal hypothalamus

**DOI:** 10.1101/2022.08.31.506043

**Authors:** Kavitha Chinnaiya, Sarah Burbridge, Aragorn Jones, Dong Won Kim, Elsie Place, Elizabeth Manning, Ian Groves, Changyu Sun, Matthew Towers, Seth Blackshaw, Marysia Placzek

## Abstract

The tuberal hypothalamus houses several major hypothalamic nuclei, dozens of transcriptionally distinct cell types, and clinically relevant cell populations implicated in obesity and related metabolic disorders. Building on recent advances in the field, here we draw upon transcriptional, signalling, and fate mapping analyses of chicken embryos and neuroepithelial explants to analyze tuberal hypothalamic development. We show that a wave of BMP signalling sweeps through early floor plate-like progenitors overlying prospective Rathke’s pouch as they track anteriorly. The timing of BMP signalling correlates with cell fate, with anterior tuberal specification complete by Hamilton-Hamburger (HH) stage 10 but posterior tuberal progenitors requiring BMPs after this point. scRNA-Seq profiling of *FGF10*-expressing cells, a proxy for cells with active BMP signalling, through HH8-21 reveals transcriptional differences that may underlie their differing response to BMPs, and the switch from neuroepithelial progenitors to stem-like radial glial cells. This study provides an integrated account of the development of the tuberal hypothalamus.

## Introduction

The hypothalamus constitutes the ventral-most part of the forebrain, and, via its links to other brain regions and to the pituitary gland, centrally regulates a broad range of behaviours and homeostatic physiological processes essential to life (Saper and Lowell, 2014). In recent years, great strides have been made in classifying the hypothalamic neurons that govern these physiological functions. ScRNA-Seq analyses and large-scale expression studies in mouse have profiled and spatially mapped major hypothalamic neuronal subtypes (Kim et al., 2022, Lee et al., 2018, Romanov et al., 2020, Shimogori et al., 2010), and in combination with classic genetic studies (Aslanpour et al., 2020, Bedont et al., 2014, Chen et al., 2020, Liu et al., 2015, Lu et al., 2013, Newman et al., 2018a, Newman et al., 2018b, Pak et al., 2014, Salvatierra et al., 2014) have identified key molecular regulators of hypothalamic neurogenesis and gliogenesis. More recently, scRNA-Seq studies covering early hypothalamic neurogenesis in embryonic chick, mouse, and human have identified markers of the major hypothalamic progenitor subtypes, and provided insights into the molecular mechanisms controlling their specification (Kim et al., 2022, Kim et al., 2020, Romanov et al., 2020, Zhou et al., 2022, Zhou et al., 2020).

One such study in chick described two spatially and molecularly distinct clusters of hypothalamic progenitors: a ‘tuberal’ progenitor stream that gives rise to neurons of the tuberal hypothalamus including neurons of the ventromedial and arcuate nuclei; and a ‘mammillary’ progenitor stream that gives rise to neurons of the mammillary and supramammillary hypothalamus, and possibly also the paraventricular nucleus (Kim et al., 2022). The tuberal progenitor domain is marked by a highly conserved set of transcription factors and signalling ligands including *SIX3, SIX6, TBX2, TBX3 RAX, FGF10* and *SHH* (Corman et al., 2018, Goodman et al., 2020, Kim et al., 2022, Kim et al., 2020, Liu et al., 2013, Muthu et al., 2016, Newman et al., 2018a, Newman et al., 2018b, Orquera et al., 2016, Pontecorvi et al., 2008, Ratié et al., 2013, Shimogori et al., 2010, Zhou et al., 2020, Brinkmeier et al., 2007, Ferran et al., 2015, Fu et al., 2017, Jeong et al., 2008, Ware et al., 2016). Following the onset of neurogenesis, tuberal progenitor subsets are apparent: anterior tuberal progenitors express higher levels of *SIX6/SIX3/SHH* while posterior tuberal progenitors express *TBX2/TBX3/FGF10* and high *RAX* (Kim et al., 2022). Pharmacological and genetic studies indicate that the specification of tuberal progenitors is governed by a diverse, evolutionarily-conserved set of cues, including the BMP, SHH, WNT, and Notch signalling pathways. BMPs and low WNT levels direct the specification of posterior tuberal progenitors, through upregulation of *TBX2/FGF10; TBX2/TBX3* in turn downregulate *SHH* (Manning et al., 2006; Trowe et al., 2013). SHH and Notch signalling, by contrast, direct anterior tuberal progenitor growth and neurogenesis (Blaess et al., 2014, Carreno et al., 2017, Corman et al., 2018, Dupé et al., 2011, Fu et al., 2017, Ratié et al., 2013, Shimogori et al., 2010, Aujla et al., 2013, Hamdi-Rozé et al., 2020, Orquera et al., 2016, Ratié et al., 2014, Szabó et al., 2009, Ware et al., 2016). Studies in mouse furthermore suggest that a delicate balance of SHH and BMP signalling directs tuberal hypothalamic development (Corman et al., 2018).

While these, and additional studies have pinpointed key molecular players that regulate the specification of tuberal cells, we are far from having an integrated understanding of how this process unfolds in time and space. Notably, an earlier fate-mapping study demonstrated that *FGF10*-expressing hypothalamic floor platelike progenitors (previously termed rostral diencephalic ventral midline (RDVM) cells (Dale et al., 1997; Dale et al., 1999) give rise to both tuberal (previously termed anterior) progenitors as well as to *EMX2*-positive mammillary (previously termed posterior) progenitors (Fu et al., 2017). In this model, *FGF10*-expressing cells are a self-renewing population that feeds out first tuberal, and then mammillary, progenitors at its anterior and posterior edges, respectively (Fu et al., 2019, Fu et al., 2017). However, how these findings relate to other signalling pathways and newer insights from scRNA-Seq have not been explored.

Here, we build on our recent studies (Fu et al., 2017; Kim et al., 2022) to examine the spatiotemporal development of the tuberal hypothalamus. We show that tuberal progenitors emerge from anterior-most *FOXA2*-expressing hypothalamic floor platelike cells as they activate canonical BMP signalling. The expansion of the anterior tuberal domain correlates with the loss of pSMAD1/5/8 and the upregulation of *SHH* expression. Our work suggests that a steady stream of tuberal progenitors is generated over a sustained period of time from hypothalamic floor plate-like cells as they track anteriorly through a zone of active BMP signalling. Tuberal progenitor cells are therefore laid down in an anterior-to-posterior order as they translocate forwards, rapidly increasing the distance between floor plate markers and the ventral telencephalon. Explant studies suggest that by HH10, neuroepithelial-intrinsic cues are sufficient to drive tuberal patterning and initiate neurogenesis. Complemented by gain-of-function and loss-of-function experiments *in vivo* and *ex vivo*, these analyses suggest a conveyor-belt mechanism whereby the time over which *FOXA2*-expressing progenitors experience BMP signalling predicts their specification into anterior or posterior tuberal progenitors. *FGF10* expression overlaps with pSMAD1/5/8, indicating that *FGF10* is not expressed in a single, self-renewing progenitor population, but rather, is expressed in successively more posterior cells. The transcriptional profile of *FGF10*-positive tuberal hypothalamic progenitors changes over time, presumably underlying the differing competence of hypothalamic progenitors. This analysis also reveals the gradual upregulation of candidate molecular regulators of a gliogenic programme that likely initiate the transformation of tuberal progenitors to radial-glial like cells found in the late-embryonic and adult tuberal hypothalamus.

## Results

### A spatio-temporal pattern of neurogenesis in the developing tuberal hypothalamus

Our aggregated scRNA-Seq datasets, interrogated using RNA velocity, revealed a distinctive differentiation trajectory for tuberal hypothalamic neurogenesis that begins in *FOXA2*^(low)^/*SIX6/RAX/FGF10* progenitor cells, passes through *ASCL1/ISL1/SIX6* neurogenic precursors, and terminates in *NR5A1/POMC/SIX6* neurons (Kim et al., 2022). This study suggested a rapid explosion of the tuberal hypothalamus that was not studied in depth. We therefore profiled the expression patterns of progenitor and neurogenic markers in detail, through multiplex hybridization chain reaction (HCR) *in situ* analyses of sagittal sections, transverse sections, wholemount ventral views and hemi-views (Supplementary Figure 1A-C). Hypothalamic floor plate-like cells, defined through *FOXA2/NKX2-1/SHH* expression, are first detected at HH6-7 (Kim et al., 2022, Supplementary Figure 1D), but tuberal progenitors (expressing *SIX6/NKX2-1/SHH/FOXA2^(low)^*) only arise between HH8 and HH10 (Figure 1A, B; Supplementary Figure 1, Kim et al., 2022). Tuberal progenitors lie posterior to *FOXG1*-positive telencephalic cells, and extend through a flat portion of anterior ventral neuroepithelium (Figure 1B; Supplementary Figure 1F, G, yellow arrows) into a characteristic fold that separates tuberal progenitors from hypothalamic floor plate-like progenitors (Figure 1B; Supplementary Figure 1F, G white arrows). Thereafter, the *SIX6*-expressing tuberal hypothalamus rapidly lengthens, increasing the separation between the telencephalon and the hypothalamic floor plate, and *SIX6* expression becomes graded, with highest expression anteriorly (Figure 1C-E). Analysis of *SIX6* expression in conjunction with *DBX1* - expressed in hypothalamic floor plate and subsequently, mammillary hypothalamus - likewise shows the appearance and dramatic expansion of the tuberal hypothalamus between HH10 and HH20 (Figure 1F-J). The appearance and lengthening of the tuberal progenitor region confirms a previous study (Fu et al., 2017) and is summarised schematically in Figure 1K.

**Figure 1:**
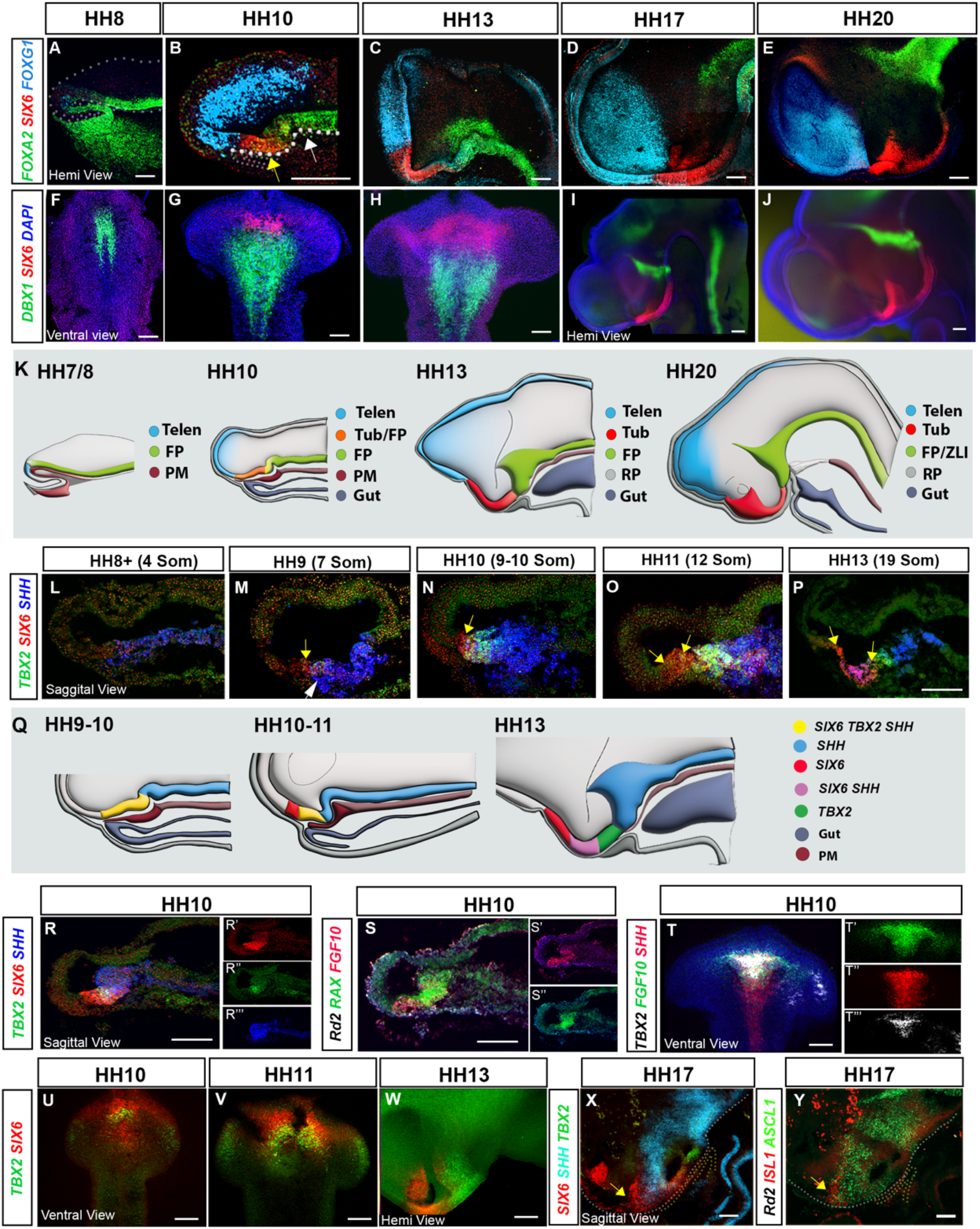
Spatio-temporal development of the tuberal hypothalamus. (A-E) Maximum intensity projections of hemi-dissected HH8-HH20 heads (‘hemi-views’) after HCR for *FOXA2, SIX6*, and *FOXG1* (yellow arrows show *SIX6/FOXA2^(low)^* region, white arrows show *FOXA2^(high)^* region). (F-J) Maximum intensity projections of isolated neuroectoderm (‘ventral views’) or hemi-views of HH8-HH20 embryos after HCR for *DBX1* and *SIX6*. (K) Schematic showing hemi-views of HH7/8-HH20; hypothalamic progenitors are shown with respect to the telencephalon and floor plate. (L-P) Sagittal sections of HH8-HH13 embryos after HCR for *TBX2, SIX6* and *SHH*. Yellow arrows indicate the extent of *SIX6*-positive, (Nagano et al., 2006)*TBX2*-negative progenitors. White arrow in M points to *SHH*-positive prechordal mesoderm. (Q) Schematic showing hemi-views of HH9-HH13 embryos, different colours represent the resolution of anterior and posterior hypothalamic progenitor domains, and floor plate. (R) Sagittal section after HCR for *TBX2, SIX6* and *SHH* at HH10. (S) Same section reprobed for *RAX* and *FGF10*. (R’-R’”; S’, S”) Single channel views of R and S. (T) Ventral view after HCR for *SHH, TBX2* and *FGF10*. (T’-T’’’) Single channel views of T. (U-W) Ventral and hemi-views after HCR for *TBX2* and *SIX6* in HH10-HH13 neuroectoderm. (X) Sagittal sections after HCR for *SIX6, SHH*, and *TBX2* and (Y) same section reprobed for neurogenic markers *ASCLl1* and *ISL1*. Yellow arrows indicate *SIX6* and *ISL1*-positive cells). All scale bars = 100 μm. Each panel shows a representative image from n = 3-5 embryos.

We next analyzed the expression of *SIX6* together with *TBX2*. At HH17, anterior tuberal progenitors are characterised by *SIX6^(high)^*-expression, and posterior tuberal progenitors, which overlie the tip of Rathke’s pouch, are characterised by *TBX2/SIX6^(low)^* expression (Kim et al., 2022; Manning et al., 2006). We investigated whether these discrete domains are already detectable at HH8, simultaneously examining their overlap with *SHH* expression.

*TBX2/SIX6/SHH*-expressing progenitors are detected between HH8-HH9 (Figure 1L, M; Supplementary Figure 1H-M), overlying the anterior-most *SHH*-expressing prechordal mesoderm (Figure 1, white arrow). These three genes are largely co-expressed, although a tiny population of *TBX2*-negative, *SIX6*-positive anterior tuberal progenitors may possibly be present (Figure 1M, yellow arrow). However, from HH10-HH13, anterior progenitors expressing *SIX6* but not *TBX2* become obvious, and expand (Figure 1N-P yellow arrows); *SHH* is expressed in posterior *SIX6*-positive cells (magenta in Figure 1P). At the same time, *SHH* starts to be downregulated in *TBX2*-expressing posterior progenitors (Figure 1N-P; expression profiles are summarised in Figure 1Q). Tuberal markers are detected in anterior-most parts of a broader *FGF10/RAX-positive* domain at HH10 (Figure 1R-T), as implied by both scRNA-seq (Kim et al., 2022) and fate mapping analysis (Fu et al., 2017). Wholemount and hemi-views clearly reveal the initial overlap, and subsequent resolution of *SIX6*-expressing anterior and *TBX2*-expressing posterior progenitors (Fig 1U-W). Multiplex HCR analysis at HH17 confirms that neurogenic (*ASCL1*-positive) cells are found within the *SIX6/SHH*-expressing region, just posterior to more differentiated tuberal neuronal precursor cells expressing *SIX6/ISL1* (Figure 1X, Y).

In summary, tuberal progenitors are first detected at HH8-9, a few hours after the appearance of hypothalamic floor plate-like cells at HH6-7, and constitute the anterior-most hypothalamic cell type. At HH8-9 tuberal progenitors have a mixed marker profile (*SIX6/TBX2/SHH/FOXA2^(low)^*), but by fine degrees these resolve to label *SIX6* and *SIX6/SHH* anterior tuberal progenitors, *TBX2* posterior tuberal progenitors, and *FOXA2*^(high)^/*SHH*-expressing hypothalamic floor plate cells at HH13. By HH17, neurogenic and more differentiated neuronal markers are detected in anterior tuberal progenitors. The spatio-temporal resolution of markers therefore predicts the spatio-temporal pattern of tuberal neurogenesis.

### BMP signalling is activated in a wave-front of hypothalamic floor plate cells as the tuberal hypothalamus is progressively generated

*TBX2* expression is regulated by BMPs in the developing tuberal hypothalamus (Manning et al., 2006) prompting us to investigate canonical BMP signalling. We detect phosphorylated (p)SMAD1/5/8 at the same time as tuberal progenitors are first specified between HH8-HH9, and pSmad1/5/8 co-localises with *FOXA2*^(low)^-expressing nuclei (Figure 2A-B”’). Thereafter, and at least until HH17 (the latest stage analysed) pSMAD1/5/8 is detected specifically in *FOXA2*^(low)^-expressing cells (Figure 2C-C”’). Analysis of pSMAD1/5/8 in combination with a range of regionspecific markers, over the period HH10-HH17, confirms that pSMAD1/5/8-positive cells overlap with, but extend posteriorly to, all other tuberal markers, including *TBX2* (Supplementary Figure 2). At all times therefore pSMAD1/5/8 is detected in posterior-most cells of the lengthening tuberal hypothalamus. At all times, pSmad1/5/8-positive nuclei lie within/overlap with the *FGF10*-expressing domain (Supplementary Figure 2B and see Figures 3O, Q), suggesting that at least a subset of *FGF10*-expressing progenitors show active BMP signalling.

**Figure 2:**
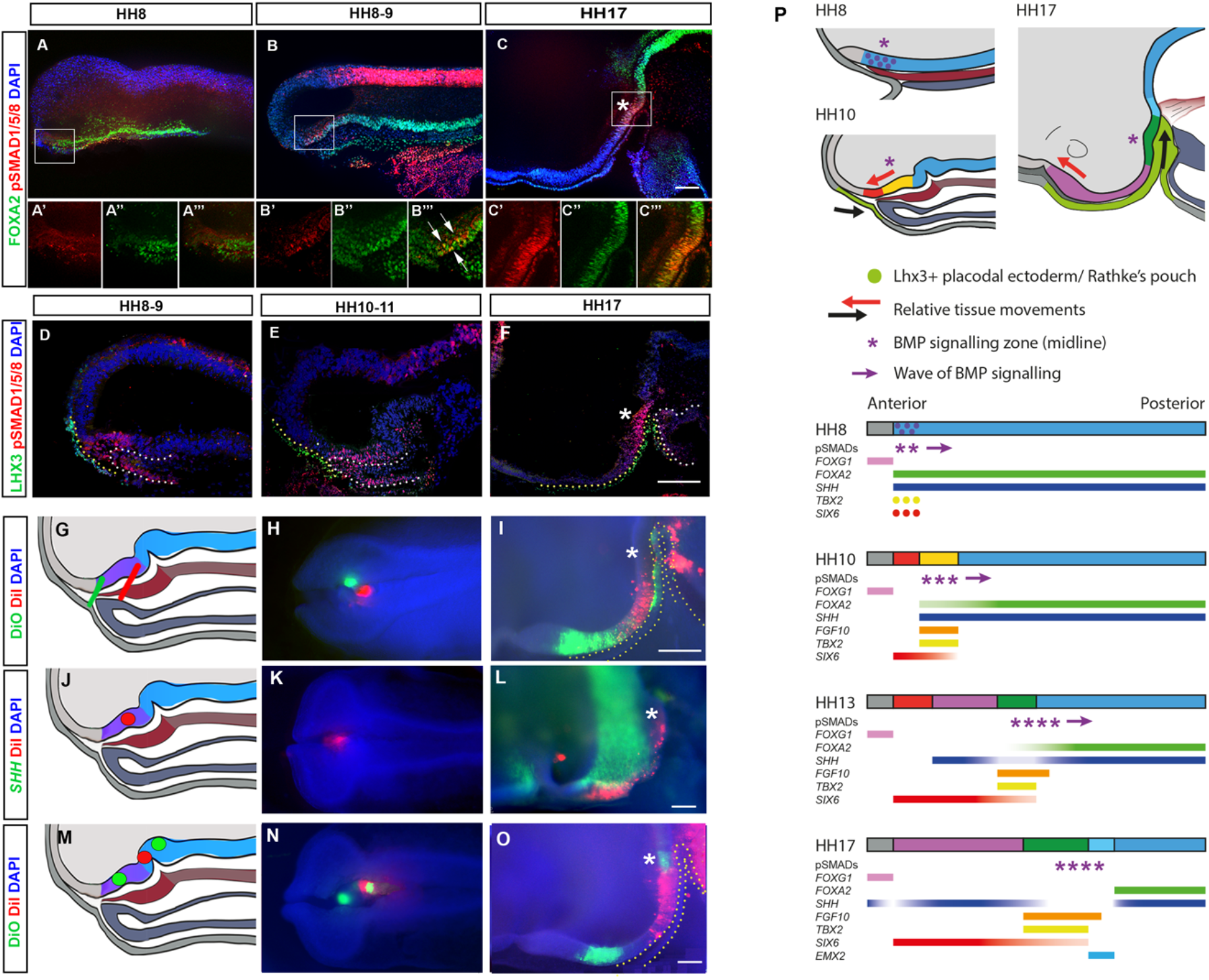
Tuberal progenitors originate from floor plate-like cells in a conveyorbelt manner. (A-C) Maximum intensity projections of hemi-dissected heads at HH8 (A) HH8-9 (B) and HH17 (C), immunolabelled for pSMAD1/5/8 and FOXA2. (A’-C’’’) High magnification single channel views of boxed regions in A-C. White arrows point to double-labelled nuclei. (D-F) Sagittal sections, immunolabelled for LHX3 and pSMAD1/5/8 in HH8-9 (D) HH10-11 (E) and HH17 (F) embryos. Yellow dots outline surface ectoderm and white dots outline the anterior gut. (G, J, M) Schematic representation of DiI/DiO injection sites shown in (H, K, N), respectively. Purple area in schematics depicts pSMAD1/5/8-positive domain). (I) Embryo shown in G, H developed to HH17 showing DiO in the anterior tuberal progenitor region and Rathke’s pouch, and DiI in the posterior tuberal progenitor region and the developing gut. (L) Embryo shown in L, K developed to HH17 and labelled for *SHH* by HCR. DiI is located in the SHH-positive anterior tuberal progenitor domain, petering out in the SHH-negative posterior tuberal progenitor domain. (O) Embryo shown in M, N developed to HH17 showing DiO in the anterior tuberal hypothalamus and (from a second injection) at the site of pSMAD1/5/8-positive cells, and DiI in the posterior tuberal hypothalamus and gut. Asterisk * indicates location of pSMAD1/5/8 cells. (P) Schematics show tissue movements as the tuberal hypothalamus expands (top) and BMP signalling and gene expression in the neuroectoderm (bottom; colour coding as per upper part and Figure 1Q). At HH8 *FOXA2* is expressed in the anterior ventral neuroectoderm. At HH10 *FOXA2* is downregulated in cells dorsal to the prechordal mesoderm (PM), which now show active BMP signalling (pSMAD1/5/8-positive nuclei), *TBX2, SIX6*, and *FGF10*. At HH13 and HH17, pSmad1/5/8 tracks back through the ventral neuroectoderm, *SHH* is secondarily upregulated in *SIX6*-expressing tissue, and the tuberal domain expands rapidly. Red and black arrows show opposite tissue movements of the neuroectoderm versus the LHX3-positive placodal ectoderm/Rathke’s pouch, respectively. Purple arrows indicate a wave of BMP signalling tracking posteriorly through the neuroectoderm. All scale bars = 100 μm: Each panel shows a representative image from n = 4 embryos/stage for immunolabelling, and from n >15 embryos for fate-mapping.

Throughout HH9-HH17, pSMAD1/5/8-positive posterior tuberal progenitors lie close to LHX3-expressing oral ectoderm/Rathke’s pouch tip cells and to pSMAD1/5/8-positive prechordal mesendoderm/anterior gut (Figure 2D-F). This raises the possibility that these are stable cell populations (at least between HH9 and HH17) that maintain their same relative positions. In this case, anterior tuberal progenitors would at all times be generated anterior to pSMAD1/5/8-positive cells. We tested this idea by DiI/DiO labelling the pSMAD1/5/8-positive domain, and simultaneously labelling cells in underlying germ layers at HH10 (Figure 2G, H; Supplementary Figure 3A-C). Examination of embryos at HH17 showed that tissues that had been labelled together at HH10 had moved out of register, the neuroectoderm being displaced relatively forwards, and Rathke’s pouch and anterior gut relatively backward (Figure 2I). DiO/Dil-labelled cells originating in the pSMAD1/5/8-positive region at HH10 (purple area in Figure 2G) were now located in the domain occupied by anterior tuberal progenitors at HH17, anterior to the pSMAD1/5/8-positive tuberal population (asterisks in Figure 2F, I), and were now pSMAD1/5/8-negative. Similar analyses, in which the pSMAD1/5/8-positive region was labelled at HH10, and then HH17 embryos analysed with *SHH* at HH17 confirmed this conclusion (Figure 2J-L). This implies that *SIX6/SHH/ASCL1/ISL1* anterior tuberal neurogenic progenitors are derived from *TBX2*-positive progenitors that transiently activate BMP signalling.

We next targeted hypothalamic floor plate-like cells posterior to the flexure, which occupy a *FOXA2^(high)^/TBX2*-positive, pSMAD1/5/8-negative domain at HH10 (Figure 2M, N, posterior DiO spot). These translocate to the HH17 region occupied by posterior tuberal progenitors (as judged through position and expression of *FGF10/SIX6^(low)^*), sometimes even just extending into the region occupied by anterior tuberal progenitors (*SIX6^(high)^/FGF10*-negative (Figure 2O; Supplementary Figure 2D-H). Triple label analyses show that cells maintain their relative spatial positions within the neuroectoderm between HH10 and HH17, with no evidence of mixing or dispersal (Figure 2M-O).

These results suggest that the tuberal hypothalamus is generated through a conveyor-belt mechanism, through a programme that is initiated when canonical BMP signalling is activated in anterior-most *FOXA2*-expressing hypothalamic floor plate-like cells (schematised in Figure 2P). The appearance of pSMAD1/5/8-positive nuclei is associated with the loss of *FOXA2* and upregulation of tuberal progenitor markers. As the neuroectoderm and underlying tissues slide past each other, prospective tuberal cells move anteriorly with respect to both underlying tissues and the zone of active BMP signalling. Posterior tuberal markers remain closely associated with the pSmad1/5/8-positive zone, while the anterior tuberal domain expands as more cells feed out of this region. Hence, prospective anterior tuberal cells move through the BMP signalling zone earlier than prospective posterior tuberal cells, suggesting that tuberal progenitor cells are generated sequentially.

### A neuroepithelial-intrinsic programme drives tuberal hypothalamic development from HH10 onward

From HH9-HH17, the LHX3-positive surface ectoderm, and later the tip of Rathke’s pouch express *BMP2* and *BMP7* (Figure 3A-F, arrows). This raises the possibility that Rathke’s pouch elicits BMP signalling in hypothalamic floor plate-like cells, the posterior-directed growth of Rathke’s pouch driving the anterior-to-posterior wave of transient pSMAD1/5/8. However, *BMP2* and *BMP7* are also expressed in hypothalamic cells (Figure 3A-F; Kim et al., 2022). Indeed, HCR analysis shows that *TBX2* is first upregulated in cells that express high levels of *BMP2* and lie just anterior to *BMP7*-expressing cells (Supplementary Figure 4A-C). Neuroepithelial-intrinsic BMPs could therefore drive tuberal development independently of signals from Rathke’s pouch and/or other extrinsic tissues such as the PM, previously implicated in hypothalamic induction (Dale et al., 1997; Dale et al., 1999). To distinguish these possibilities, we isolated prospective tuberal progenitor cells from other tissues at HH10 (Figure 3G-G’’’’’). The accuracy of dissection was confirmed by examining the molecular profile of acutely-dissected explants, which, as *in vivo* (Figure 3H, I; Supplementary Figure 5A-D), are positive for pSMAD1/5/8, *SHH* and *FGF10*, but not *FOXG1, ISL1*, or *EMX2* (Figure 3J-K”; Supplementary Figure 5E-H). Explants cultured to a HH13 equivalent grew, and generated domains that showed the same organisation as *in vivo. SHH* and *FGF10*, which are co-expressed throughout the HH10 explant (Figure 3K-K”), resolved into overlapping domains (Figure 3L-L”), and distinct but overlapping regions expressing *SIX6, SIX6/SHH*, and *TBX2* were detected, as *in vivo* (compare Figure 3M-M” to Figure 1P). After culture to a HH24 equivalent, explants had grown extensively and showed discrete but overlapping domains of gene expression, composed of *ISL1/SHH* anterior tuberal neurogenic progenitors, adjacent to pSMAD1/5/8-positive posterior tuberal progenitors, and then *EMX2*-expressing mammillary progenitors (Figure 3N-N’’’’). Isolated explants, free from any possible influence of surrounding tissues, therefore grow to generate a spatial pattern that is similar to that *in vivo* (Figure 3O-Q). We conclude that, although BMP ligands from non-hypothalamic structures are required to initiate tuberal development (Dale et al., 1997, Dale et al., 1999), from HH10 onwards, hypothalamic neuroepithelium-intrinsic factors are sufficient to maintain tuberal hypothalamic regionalization and neurogenesis.

**Figure 3:**
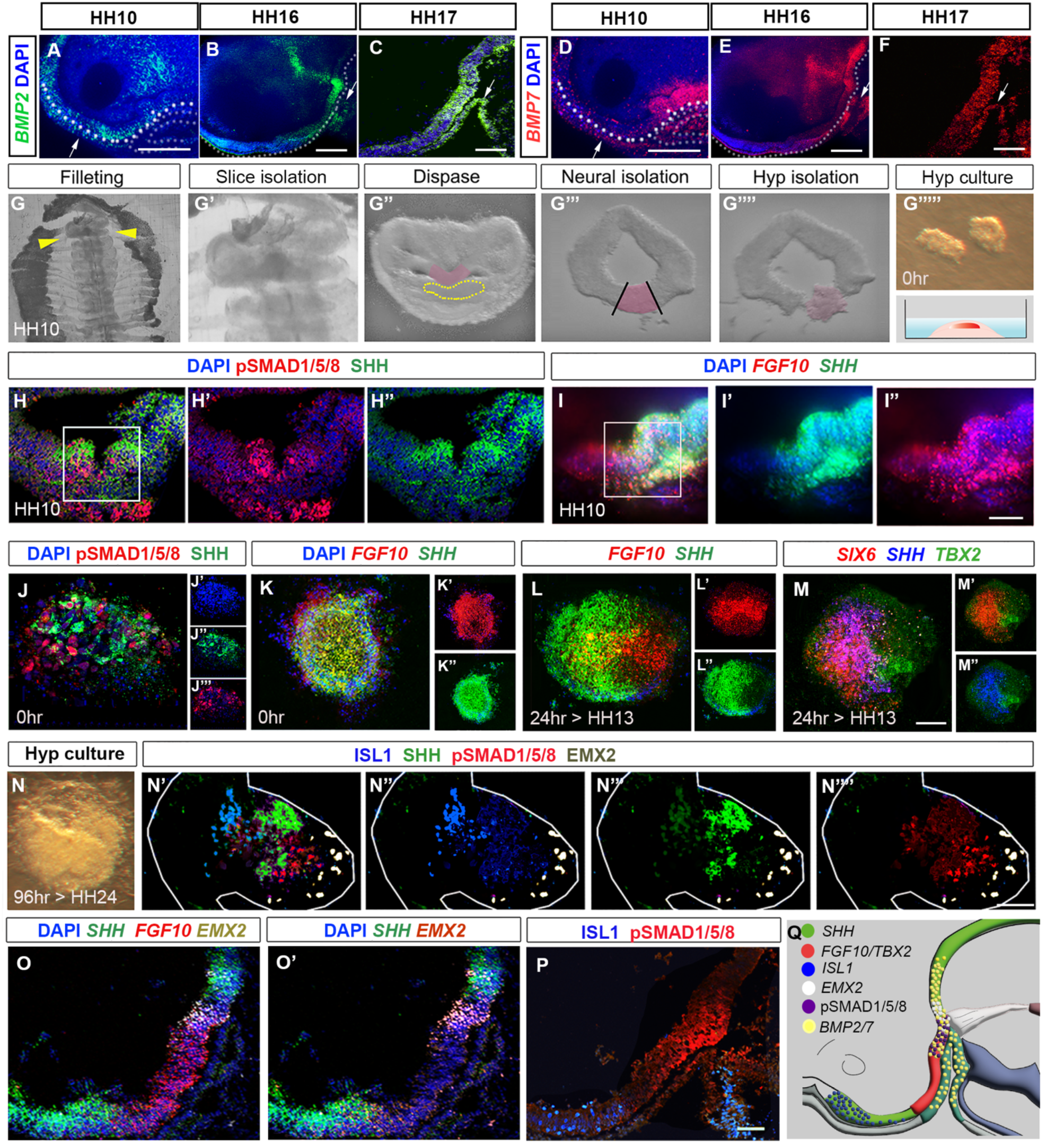
Organisation of tuberal domains through a neuroepithelial-intrinsic mechanism. (A-F) Maximum intensity projections of hemi-dissected heads after HCR, showing expression of *BMP2* or *BMP7* at HH10, HH16 and HH17. (G) Steps describing hypothalamic tissue isolation from a HH10 embryo: (G, G’) filleting, yellow arrows in G mark the position of the slice containing hypothalamic tissue, shown at higher power in G’. (G”) Isolated slices containing hypothalamic tissue have a characteristic shape, and the prechordal mesoderm can be identified morphologically (dotted outline). (G”’) After dispase treatment, neural tissue is isolated from surrounding tissues, and hypothalamic tissue (pink) excised (G””). (G’’’’’) Isolated hypothalamic tissue is embedded in a 3-D collagen matrix. (H-I’) Sagittal sections at HH10, immunolabelled to detect pSMAD1/5/8 and SHH (H: box marks enlarged area shown as single channel views in H’, H”), or analysed by HCR to detect *SHH* and *FGF10* (I: box marks enlarged area shown as single channel views in I’ and I’’). (J-N””) Sections through hypothalamic explants, at 0hr, or cultured for 24hr or 80hr, analysed by immunolabelling or HCR. (J-J”) At 0hr, pSMAD1/5/8 and SHH are detected throughout the section (single channel views shown in J’-J”’-blue shows DAPI counterstain). (K-K”) At 0hr, *FGF10* and *SHH* are detected throughout the section (single channel views shown in K’, K”). (L-M”) Ater a 24hr culture period, to the equivalent of HH13, *FGF10* and *SHH* begin to resolve (L: single channel views shown in L’, L”), and discrete domains of *SIX6, SHH*, and *TBX2* are apparent (M: single channel views shown in M’, M”). (N) Brightfield image of explant cultured for 96 hrs to the equivalent of HH19. (N’-N’’’’) Organised expression of *ISL1, SHH*, pSMAD1/5/8, and *EMX2* at 96 hrs. (O-P) Serial adjacent sagittal sections of HH19 embryos, analysed by HCR to show expression of *SHH, FGF10* and *EMX2* (O; O’ shows same section without *FGF10)*, or by immunolabelling to detect ISL1 and pSMAD1/5/8 (P). (Q) Schematic summarising the expression of *SHH, TBX2, FGF10, ISL1, EMX2*, pSMAD1/5/8 and *BMP2/7* at HH24. Scale bars = 100uM. Each panel shows a representative image from a minimum of n= 3 embryos or explants.

### BMP signalling directs tuberal development during HH9-HH13

Our findings predict that exposure of hypothalamic floor plate-like cells to BMPs will lead to the generation of tuberal cells. To test this, we performed *ex vivo* studies, building on previous work that has delineated the spatial position of future hypothalamic regions and the time at which they are specified (Kim et al., 2022, Ohyama et al., 2005). We isolated ‘anterior’ *NKX2-1*-positive neural explants that fate-map to the tuberal hypothalamus, and *NKX2-1*-positive ‘posterior’ neural explants, that fate-map to the mammillary hypothalamus (Figure 4A, D). After culture to a ~HH14 equivalent, anterior control explants expressed tuberal progenitor markers (*SIX6/NKX2.1/SHH/TBX2)*, and anterior-tuberal neuronal markers (*NR5A1, POMC)* (Figure 4B-B’’’’’’). By contrast, posterior control explants expressed *SHH/FOXA2* (Figure 4C’’’, E’’, E”’), markers that define the later floor plate/supramammillary hypothalamus (Kim et al., 2022). However, posterior explants that were exposed to 32nM BMP2/7 upregulated the tuberal markers *TBX2*, *SIX6*, and *NR5A1* (Figure 4F’, F’’’’, F’’’’’), and after a more prolonged culture to a HH18 equivalent, *POMC* (Figure 4G, H) (note that *in vivo, POMC* is detected only after *NR5A1* (Kim et al., 2022). We conclude that BMP2/7 can induce hypothalamic floor plate-like cells to form tuberal cells.

**Figure 4:**
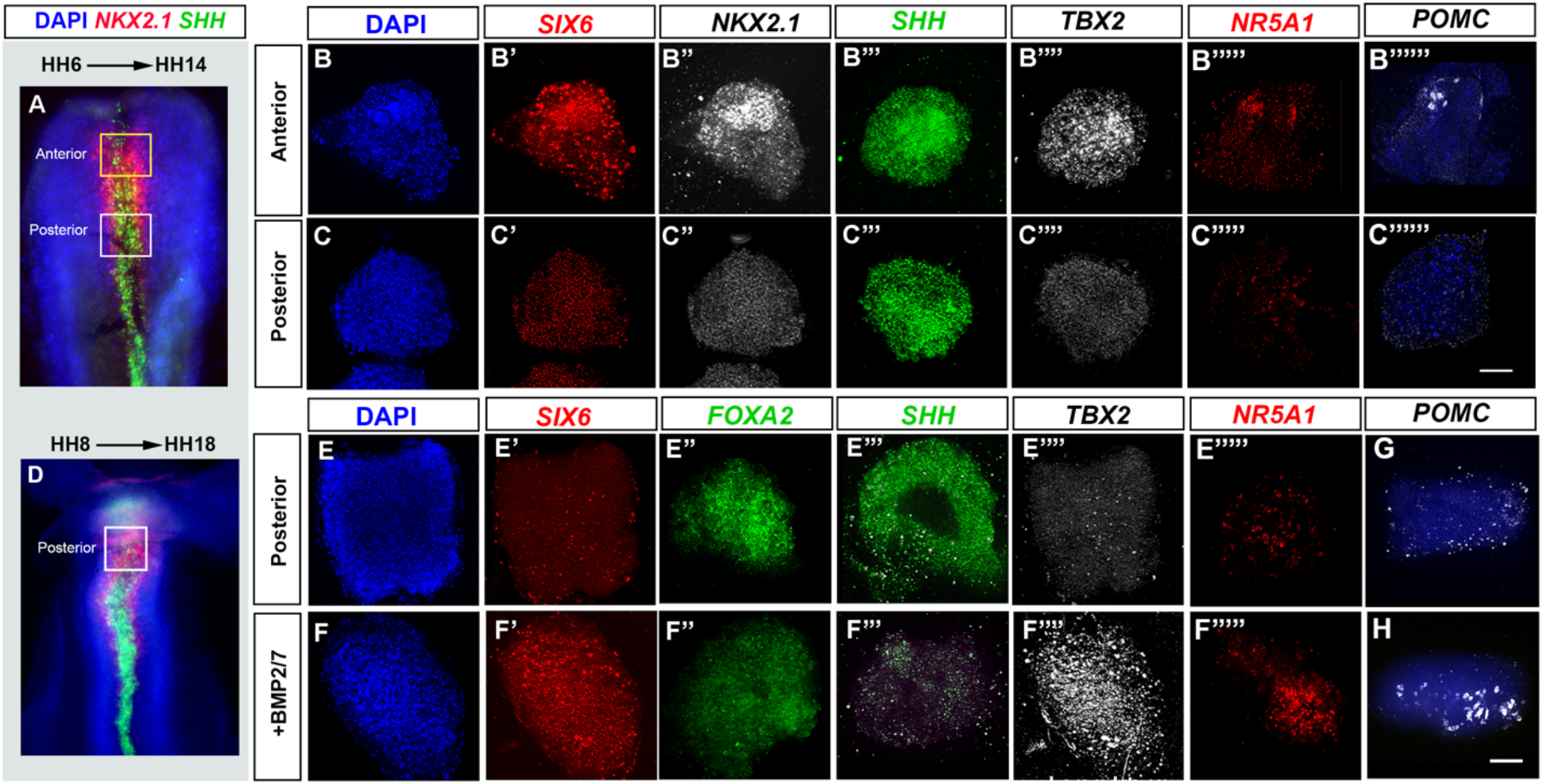
BMP promotes tuberal identity in hypothalamic floor plate-like cells. (A, D) HH6 (A) or HH8 (D) isolated neuroepithelia, after HCR to detect *NKX2-1* and *SHH*. Boxes show regions dissected for explant culture. (B-B”””, C-C”””) HH6 anterior (row B) and posterior (row C) explants cultured to a HH14 equivalent and processed by wholemount HCR for *SIX6, NKX2.1, SHH, TBX2, NR5A1 AND POMC*. (E-F’’’’’) HH6 posterior explants cultured to a HH14 equivalent in control medium (row E) or medium containing 32 nM BMP2/7 protein (row F) and processed by wholemount HCR for *SIX6, FOXA2, SHH, TBX2*, and *NR5A1*. (G-H) HH8 posterior explants cultured to a HH18 equivalent and processed by wholemount HCR for *POMC*. Scale bars = 100μm. Each panel shows a representative image; *NKX2-1, NR5A1, POMC, FOXA2* were analysed in a minimum of n=3 explants/condition; *SIX6, SHH, TBX2* were analysed in a minimum of n=11 HH6 explants/condition and n=7 HH8 explants/condition.

To complement these studies, we tested whether a reduction in BMP signalling suppresses tuberal hypothalamic development in vivo. Implantation of beads, soaked with the BMP inhibitor Noggin (Smith and Harland, 1992) in the tuberal region of a HH9-HH10 embryo (Figure 5A) resulted in an acute - albeit transient - loss of pSMAD1/5/8 (Supplementary Figure 6). This resulted in partial suppression of tuberal development at HH18 (Figure 5B-C’). The area occupied by *SIX6*-expressing anterior tuberal progenitors was significantly reduced, there was no/little sign of the *TBX2*-positive posterior tuberal progenitor domain, and *SHH* was abnormally maintained in the ventral region where *TBX2* is normally expressed (Figure 5B-D).

**Figure 5:**
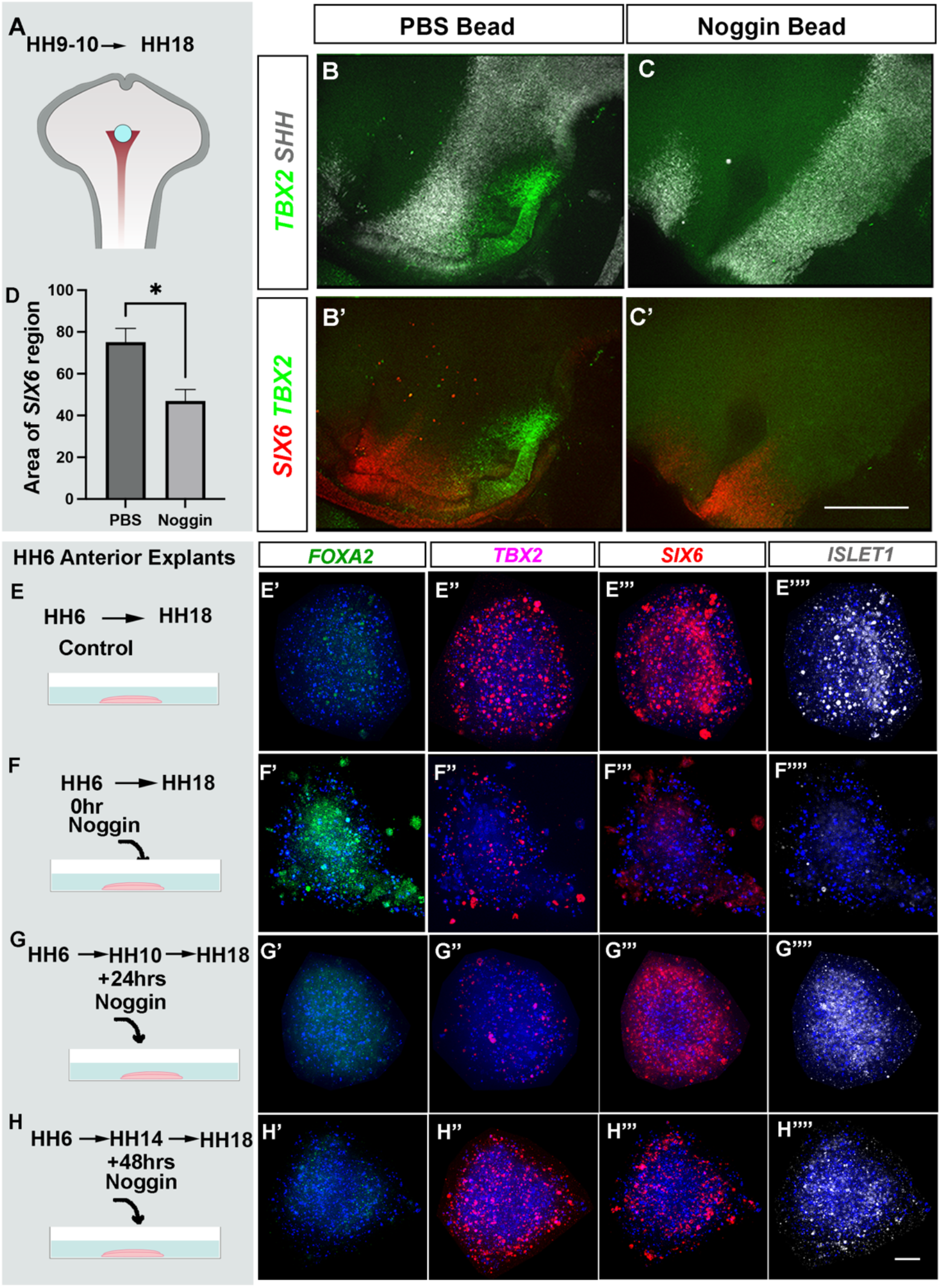
BMP signalling temporally patterns anterior and posterior tuberal progenitors. (A) Schematic depicting position of bead implantation at HH10. (B-C’) Hemi-views of HH18 heads, analysed by triple label HCR to detect *TBX2, SHH* and *SIX6* after PBS (B, B’) or Noggin (C, C’) bead implantation. (B, C) show *TBX2* in combination with *SHH;* (B’,C’) show *TBX2* in combination with *SIX6*. (D) There is a significant decrease in the area of the *SIX6*-positive domain after exposure to Noggin (*P < 0.05\**, unpaired t-test). n= 5 embryos/condition. (E-H) Schematics show experimental design to examine the impact of Noggin exposure at different times to tuberal specification. HH6 anterior explants, cultured for 72 hrs to a HH18 equivalent, in the absence of Noggin (E), or (F-H) after exposure to 300 ng/ul Noggin at the onset of culture (0hr, F), after 24hr culture (G) or after 48hr culture (H). (E’-E””; F’-F””; G’-G””; H’-H””) Wholemount analyses of explants cultured to a HH18 equivalent, analysed by HCR to detect *FOXA2, TBX2, SIX6 and ISL1*. Each row shows a single explant. (n=4 explants (row E); n=3 explants (row F); n=3 explants (row G); n= 5 explants (row H). All scale bars = 100 μm.

The Noggin-mediated transient reduction of BMP signalling did not, however, completely eliminate tuberal cells. This likely reflects the fact that tuberal hypothalamic specification has already been initiated by HH6 (Figure 4A-B””), and that pSMAD1/5/8 is already detected ~6 hrs later, at HH8-9 (Figure 2B, D). We therefore sought to eliminate BMP signalling at HH6. Bead implantation is unreliable at this stage, so we again turned to the HH6 *ex vivo* assay. Anterior explants cultured in control media to the equivalent of ~HH18 did not express the supramammillary/floor plate marker, *FOXA2*, but expressed the posterior tuberal marker *TBX2* and the anterior tuberal markers *SIX6* and *ISL1* (Figure 5E-E””). In contrast, anterior explants cultured from the outset in 30nM Noggin showed little or no expression of either *SIX6, TBX2*, or *ISL1*, but instead, expressed *FOXA2* (Figure 5F-F””). To determine whether ongoing BMP signalling is needed to maintain tuberal development, we inhibited BMPs at later timepoints within the same experimental setup. We again isolated anterior explants at HH6, cultured them for 24 hrs in control medium to the equivalent of HH10, then exposed them to Noggin until the equivalent of HH18. Under these conditions, *FOXA2* was lost and the explants expressed *SIX6* and *ISL1*, but not *TBX2* (Figure 5G-G””). This is in close agreement with the *in vivo* experiment, and indicates that BMP signalling prior to HH10 is sufficient for the expression of anterior tuberal markers, but a longer or later period of BMP exposure is required for posterior tuberal identity. Finally, the addition of Noggin to anterior cultures at 48 hrs, which is the equivalent of ~HH14, had no effect on the markers studied, with *FOXA2, TBX2, SIX6*, and *ISL1* expression similar to explants cultured in the control medium (Figure 5H-H””). This indicates that BMPs are needed between HH6 and HH14 for the proper induction of posterior tuberal progenitors.

### Excessive BMP levels disrupt tuberal progenitor tissue homeostasis

Our fate mapping studies showed that *SIX6/SHH/ASCL1/ISL1* anterior tuberal neurogenic progenitors are derived from progenitors that are only transiently *FOXA2-, TBX2*-, and pSMAD1/5/8-positive. This raises the possibility that the cessation of BMP signalling is an important step in tuberal development, and we reasoned that prolonged/excessive exposure to BMPs may disrupt the tuberal programme. To directly test this, PBS-control or BMP2/7-soaked beads were implanted in anterior-most tuberal regions of HH10 embryos and embryos examined at HH13/14, each taken through two rounds of multiplex HCR. Ectopic BMPs significantly reduced, or eliminated, *SIX6/SHH* anterior tuberal progenitors (Figure 6A-A”, C-C’’: bracket in Figure 6A’; arrow in Figure 6C’). *SHH* expression was not appropriately downregulated in *TBX2*-expressing cells (bracket in Figure 6A”, C”), and was marginally reduced in the region of the anterior ID (Kim et al., 2022; Shimogori et al., 2010). Instead, this region was occupied by an expanded domain of strongly *TBX2*-expressing posterior tuberal progenitors (Figure 6A, A”, A”’, C, C”, C”’ yellow arrows; Figure 6E). No obvious changes in expression of the prethalamic marker *PAX6* were detected, but expression of *FST*, which at this stage is restricted to a domain located just dorsal to the anterior ID, was eliminated (Figure 6B, D, white arrows). We have previously shown that prethalamic-derived FST inhibits hypothalamic specification (Kim et al., 2022). Given that BMP7 is a potential target of FST, this result raises the possibility that BMP signalling is maintained in the tuberal hypothalamus in part by suppressing expression of a BMP inhibitor.

**Figure 6:**
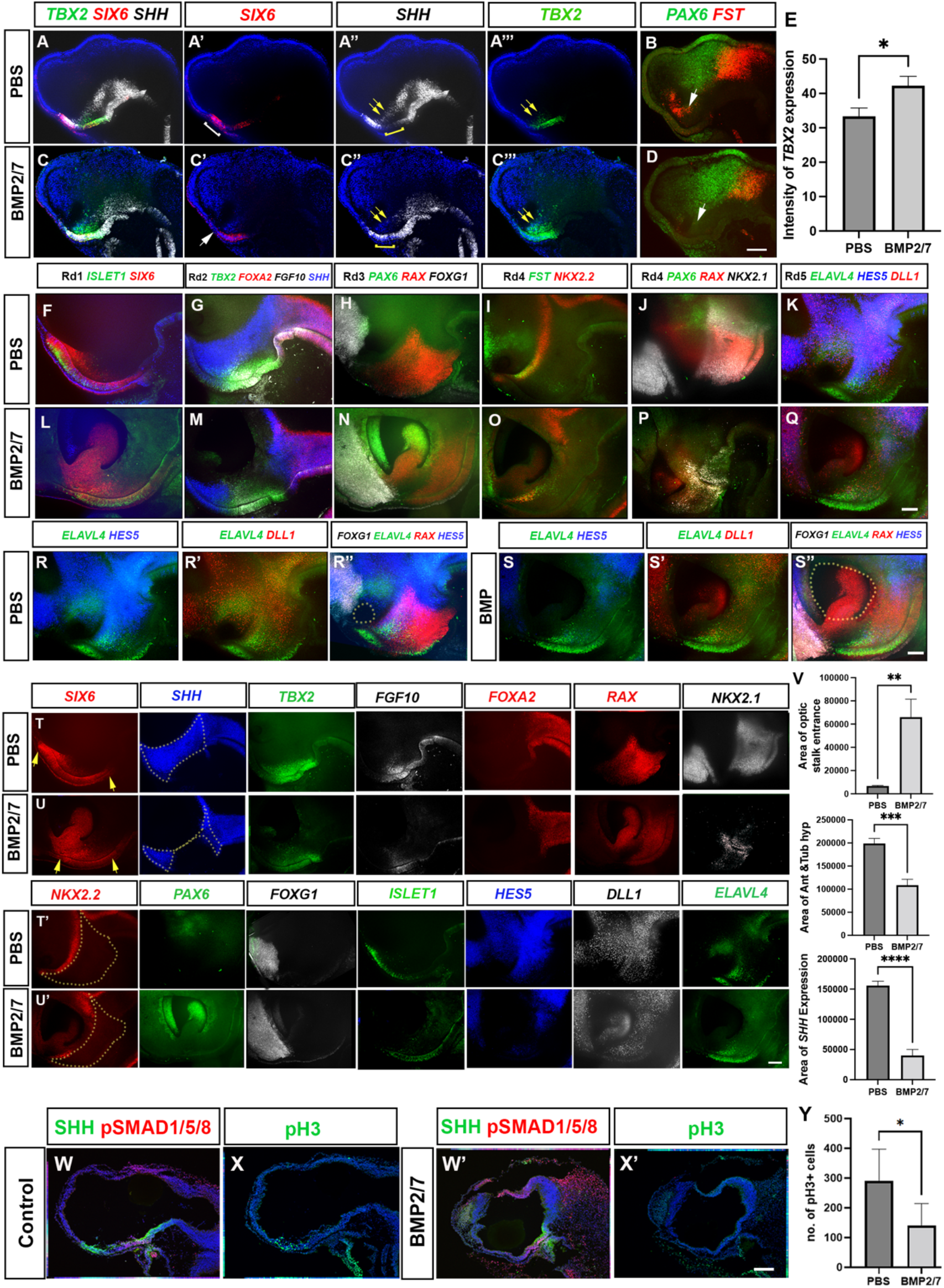
Ectopic BMP exposure reduces anterior, then posterior tuberal progenitors. (A-D) Maximum intensity projection of hemi-dissected head of HH13/14 embryos taken through 2 rounds of multiplex HCR analysis, after implantation of PBS bead (A-B) or BMP2/7-soaked bead (C-D) Round 1: analysed for expression of *TBX2, SIX6, SHH* (A-A’’’ and C-C”’). Round 2: analysed for *PAX6* and *FST* (B, D). White bracket in A’ shows *SIX6*-positive *TBX2*-negative anterior tuberal progenitors, absent after BMP2/7 exposure (white arrow in C’). Yellow bracket in A” shows *TBX2*-positive *SHH*-negative tuberal progenitors; after BMP2/7 exposure, *SHH* is unusually maintained in the midline (yellow bracket in C”), but reduced in the anterior ID (yellow arrows in A’’, C”). White arrow in B, D point to FST-expressing cells, absent after exposure to BMP2/7. (E) Graph showing a significant increase in intensity of *TBX2* expression following BMP2/7 bead implantation p<0.05, unpaired t-test n= 3 embryos/condition. (F-Q) Hemi-view of PBS and BMP2/7 bead grafted embryos incubated for 42 hr, until a HH18 equivalent (n=4 each/condition), analysed by three rounds of multiplex wholemount HCR for *SIX6, ISL1* (Round1, F, L); *SHH, TBX2, FGF10 and FOXA2* (Round 2, G, M); *PAX6, RAX* and *FOXG1* (Round 3, H, N). Embryos then subdivided and subject to one or two further rounds of multiplex wholemount HCR (n=2 embryos each for rounds 4 and 5): *FST, NKX2.2* (Round 4, I, O); or *PAX6, RAX* and *NKX2.1\** (Round 4, *different embryo, J, P); *ELAVL4, HES5*, and *DLL1* (Round 5, K, Q). (R, R’, S, S’): dual channel views from embryos shown in panels K, Q. (R”, S”) Representative images from Rounds 3 and 5 overlaid in Photoshop. Dotted outline shows optic stalk entrance. (T-U’) Individual channel views of multiplex views shown in F-Q, in PBS (T, T’) or BMP2/7-treated (U, U’) embryos. Yellow arrows point to length of *SIX6*-positive domain. Dotted outlines show areas measured. (V) Graphs show (top): a significant increase in the area of optic stalk entrance, P<0.001**, unpaired t-test (dotted lines in R’’, S’’); (middle): significant decrease in the area of anterior and tuberal hypothalamus, P<0.0004***, unpaired t-test (dotted line in T’, U’) and (bottom): a significant decrease in the area of *SHH* expression, P< 0.0001****, unpaired t-test (dotted lines in T, U *SHH* single channel view) between PBS and BMP2/7 bead-implanted embryos. (W-X’’) Serial adjacent sagittal sections of HH17 embryos, immunolabelled to detect expression of SHH, pSMAD1/5/8, or the M-phase marker phosphoH3 after implanting PBS bead (W, X) or BMP2/7-soaked bead (W’, X’) at HH10. n=6 embryos/condition (Y) Graph showing significant decrease in the number. of pH3 positive M-phase progenitors (P<0.05*, unpaired t-test). All scale bars = 100μm

A further set of embryos were examined at HH18, each taken through repeated rounds of multiplex HCR. Exposure to BMP2/7 either reduced (n=4) or eliminated (n=2) anterior tuberal progenitors, as measured through a reduction (Figure 6) or loss (Supplementary Figure 7) of *SIX6* and *SHH* expression (compare Figure 6F, G with Figure 6L, M; individual channels shown in Figure 6T, U; compare Supplementary Figure 7A, B with Supplementary Figure 7E, F; individual channels shown in Supplementary Figure 7I, J). Surprisingly, and in contrast to embryos examined at HH13, BMP-treated embryos also showed a reduction or loss of posterior tuberal progenitors, reflected in decreased (Figure 6), or absent (Supplementary Figure 7), expression of *TBX2* and *FGF10* (compare Figures 6G and M; individual channels shown in Figure 6T, U; compare Supplementary Figures 7B and F; individual channels shown in Supplementary Figure 7 I, J). The loss of tuberal progenitor domains was accompanied by marked changes in anterior ventral forebrain morphology: the optic stalk failed to narrow, resulting in a gaping entrance to the optic stalk (Figure 6V) and copious *SIX6/RAX/PAX6-positive* eye tissue within (compare Figures 6F, H, with Figures 6L, N; individual channels shown in Figure 6T,T’, U, U’; optic stalk entrance outlined in Figures 6R” versus 6S”). By contrast, the *FOXA2* expression domain, encompassing floor plate and supramammillary hypothalamus, appeared normal (compare Figures 6G and 6M; individual channels shown in Figures 6T, U; compare Supplementary Figures 7B and F; individual channels shown in Supplementary Figure 7I, J).

Additional rounds of multiplex HCR confirmed the reduction or loss of the anterior and posterior tuberal progenitor territories as well as the anterior ID, as judged by reduction or loss of expression of *RAX, NKX2-1, SHH, NKX2-2* and *FST* (compare Figures 6G–J with Figures 6M-P; individual channels shown in Figures 6T, T’, U, U’; compare Supplementary Figures 7C, D with 7G, H; individual channels shown in Supplementary Figure 7I, J). These analyses confirmed that posterior hypothalamic regions were barely affected: the supramammillary and mammillary markers *PITX2* and *EMX2* were maintained (compare Supplementary Figures 7C and 7G; individual channels shown in Supplementary Figure 7I, J). Expression of the telencephalic marker *FOXG1*, likewise, was unaffected, maintaining its relation to the optic stalk opening (compare Figures 6H, R with Figures 6N, S; individual channels shown in Figure 6 T’, U’). We conclude that excessive BMP exposure leads ultimately to a loss of expression of molecular markers of both anterior and posterior tuberal progenitors.

Quantitative analyses confirmed that the reduction in tuberal progenitor marker expression correlated with a significant decrease in the area of the tuberal hypothalamus (Figure 6V). To examine a likely mechanism for the decrease in tuberal territory, we examined proliferation in control versus BMP-treated embryos (Figure 6W-X’). BMP-treated embryos showed a significant reduction in M-phase progenitors in the tuberal hypothalamus (compare Figures 6X and X’; Figure 6Y).

In embryos in which tuberal progenitors were entirely lost after BMP2/7 exposure, *ISL1* - a marker of postmitotic tuberal neural precursors (Kim et al., 2022) - was not detected (compare Supplementary Figures 7A, I with 7E, J). However, in embryos in which tuberal progenitor markers were reduced, some *ISL1* persisted, albeit weakly (compare Figures 6F and 6L; individual channels shown in Figure 6T’, U’), In a final round of HCRs, embryos were therefore examined for expression of the Notch target gene *HES5*, the neurogenic progenitor marker *DLL1*, and the neural precursor marker *ELAVL4* (Figures 6K, Q; double channel views shown in Figures 6R, R’, S, S’). Despite their widespread expression in the brain, hypothalamic expression could be accurately determined through alignment with images from previous rounds of HCR (Figure 6R”, S”). In control embryos, *HES5, DLL1* and *ELAVL4* are all detected in the hypothalamus (Figure 6K, R-R”, T’). By contrast, *HES5* expression is dramatically reduced in BMP-exposed embryos (Figure 6Q, S-S”, U’). Notch maintains the progenitor state in chicken hypothalamus (Hamdi-Rozé et al., 2020, Place et al., 2022, Ratié et al., 2013, Ratié et al., 2014, Ware et al., 2016) so the loss of *HES5* in BMP-treated embryos may be associated with ectopic neurogenesis.

Despite this result, no clear differences in *DLL1* and *ELAVL4* expression were seen in the residual hypothalamic domain in BMP-treated embryos (Figure 6Q, U’). A likely interpretation is that an increase in neurogenesis may be balanced out by the existence of fewer progenitor cells (Figure 6X’, Y).

### Tuberal progenitors change over time

Our findings suggest that distinct spatial domains of the tuberal hypothalamus are generated sequentially as the result of transient BMP signalling. This implies that pSmad1/5/8-positive progenitors may show dynamic patterns of gene expression that control their ability to generate distinct subsets of tuberal cells. To identify such genes we digitally isolated *FGF10*-expressing cells from our HH8-HH20 scRNA-Seq dataset (Figure 7A, Kim et al., 2022) - a proxy for pSmad1/5/8-positive cells at all times (eg Supplementary Figure 2B (HH10) and compare Figures 3O and 3Q (HH19)). This analysis showed differential expression of a number of known developmental regulators between early (more anterior) and late (more posterior) *FGF10*-expressing cells (Figure 7B). *CHRD (Chordin), BMP2/7* and *SST* showed enriched expression in early-stage (HH8-10) tuberal progenitors, whereas the retinoic acid binding protein *CRABP1, WNT5A*, and the growth factor *CTGF* showed enriched expression in later-stage tuberal progenitors. The WNT/FGF pathway inhibitor *SHISA2* (Nagano et al., 2006) was expressed in early, but not late, tuberal progenitors, and the canonical WNT mediator (and adherens junction component) *CTNNB1 (β-catenin)* increased over time. Changes in expression levels of the NOTCH signalling pathway modulator *LFNG*, and its target genes (*HES5, HES5-like)* indicated a gradual increase in NOTCH signalling over time. Likewise, the transcription factors *ONECUT1, ID1, ID2, ID4, SOX8*, and *SOX9* showed enriched expression in later-stage *FGF10*-expressing tuberal progenitors. HCR analyses confirm the reduction in *BMP2/7* expression between HH8 and HH13 (Figure 3A-F), and the upregulation of *ID1, ID2, ID4* and *SOX9* in later-stage (posterior) tuberal progenitors (Figure 7C-H; Place et al 2022). In all, both abrupt (e.g. *BMP2/7, SHISA2*) and gradual changes in gene expression are detected, conferring a distinct transcriptional profile upon *FGF10*-positive cells at each stage sequenced.

**Figure 7:**
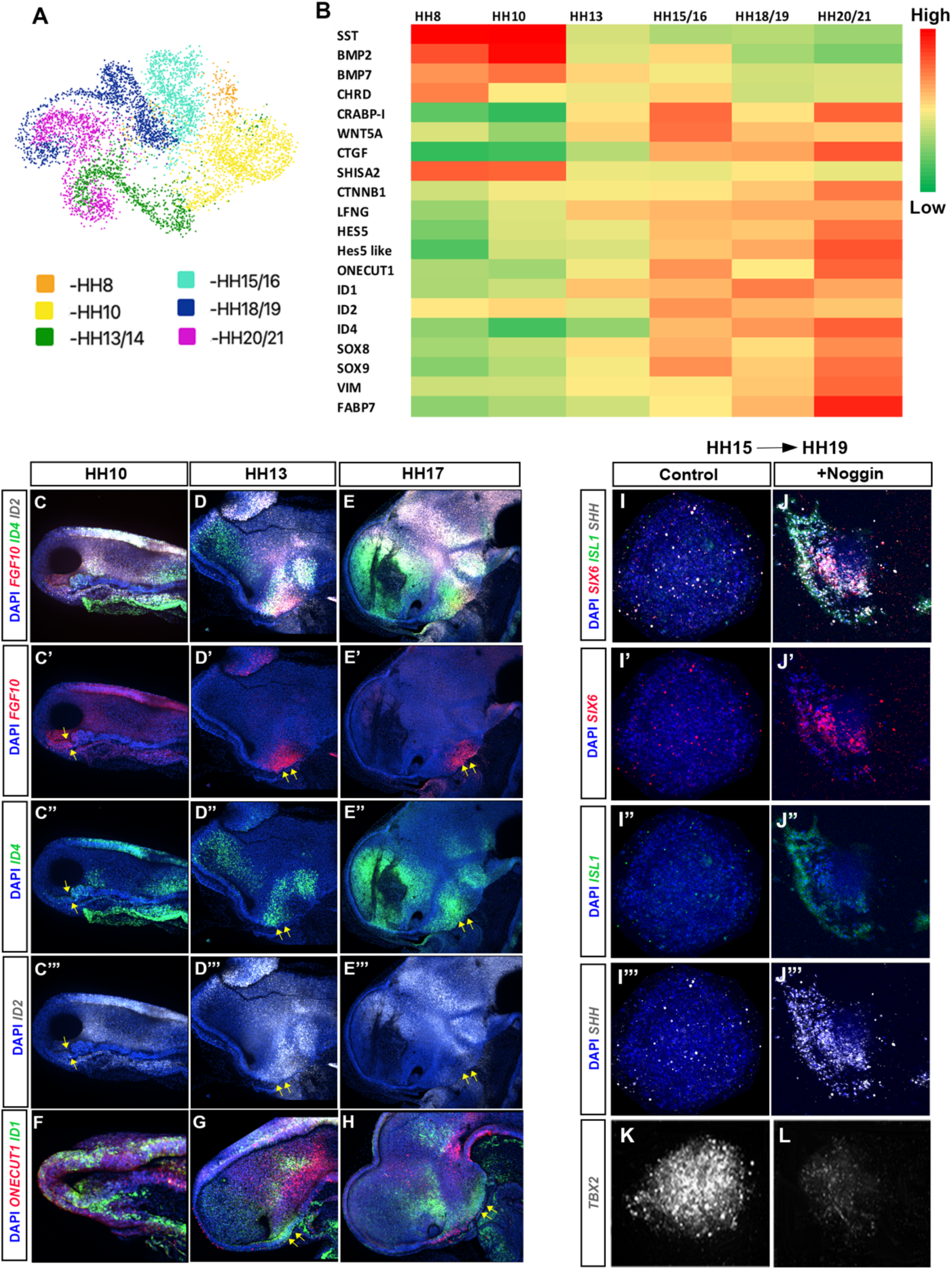
Dynamic transcriptional changes in *FGF10-expressing* tuberal progenitors. (A) UMAP plot showing the distribution of *FGF10* expressing tuberal progenitors isolated in a previous scRNA sequencing study (Kim et al., 2022). (B) Heat map showing upregulated and downregulated genes between stages HH8-HH20/21. (C-E’’’) Maximum intensity projections of hemi-dissected heads of HH10 (C), HH13 (D) and HH17 (E) embryos after multiplex HCR for *FGF10, ID4* and *ID2*. (C’-E’) Shows expression of *FGF10* with DAPI. (C”-E”) Shows expression of *ID4* and DAPI. (C’’’-E’’’) Shows expression for *ID2* and DAPI. (F-H) Maximum intensity projections of hemi dissected views of HH10 (F), HH13 (G) and HH17 (H) embryos after multiplex HCR for *ONECUT1 and ID1*. Yellow arrows point to *FGF10*-expressing domain. (I-L) Maximum intensity projections of explants dissected from the *FGF10*-expressing region at HH15 (ie region shown by arrows in (D’, E’) and cultured to a HH19 equivalent alone (I-I”’, K) or with Noggin (J-J”’, L) analysed by HCR for *SIX6, ISL1, SHH* and *TBX2*. n= 3 explants/condition analysed with *SIX6/ISL/SHH*.

Our *ex vivo* experiments had suggested that anterior and posterior tuberal progenitors are specified by HH14 (Figure 4M). Additional *ex vivo* experiments, isolating posterior tuberal progenitors at HH15, and culturing alone, or in the presence of Noggin, demonstrates that after this point, BMPs actively *repress* the anterior tuberal program (Figure 7I-L) and this switch in competence coincides with the transcriptional onset of a large set of transcription factors (Figure 7B). Upregulation of *SOX8/9* and *ID* family genes has been previously reported in latestage neuronal progenitors in mammals (Clark et al., 2019, Lyu et al., 2021, Sagner et al., 2021, Telley et al., 2019), raising the possibility that the expression of these genes in later posterior tuberal progenitors is associated with neurogenesis. Alternatively, this upregulation may be directly linked to the transition of posterior tuberal progenitors from a neuroepithelial to radial glial-like state, a process that is regulated by SOX9 in the developing spinal cord (Scott et al., 2010). In support of this, the radial glial markers *VIM* and *FABP7* are upregulated in *FGF10*-expressing progenitors at HH20/21 (Figure 7B). These molecular changes, therefore, may reflect the transition of *FGF10*-expressing posterior tuberal progenitors to the specialised, *FGF10*-expressing radial glial-like tanycytes that characterise the late-embryonic and adult tuberal hypothalamus (Goodman et al., 2020, Haan et al., 2013, Robins et al., 2013, Yoo et al., 2021).

## Discussion

Until recently, the extraordinary complexity of the hypothalamus had prevented a detailed understanding of its structure and functions. Contemporary techniques and reagents have led to an influx of new insights into its cell type composition and circuitry, at long last providing a substantial base for further discoveries (Chen et al., 2017, Kim et al., 2020, Mickelsen et al., 2020, Moffitt et al., 2018, Romanov et al., 2020, Zhou et al., 2020). Building on earlier work, we set out to explore specific mechanisms by which the tuberal hypothalamus - a region which harbours neurons and glia that govern hunger, energy balance, reproduction, response to stress, and a range of social, sexual, affiliative, and emotional behaviors (Saper & Lowell, 2014, Swaab, 2003) - is formed. We show that the tuberal hypothalamus is generated in a conveyor-belt mechanism, in which a steady stream of *FOXA2*-expressing hypothalamic floor plate cells pass through a zone of active BMP signalling, directing their specification into tuberal progenitors, and uncover candidate molecular regulators of the transition of tuberal progenitors to neurogenesis and gliogenesis.

### Specification of the tuberal neurogenic hypothalamus from hypothalamic floor plate-like cells

Many models of forebrain development suggest that forebrain cells acquire regional identity during a period where there is little growth (Finlay et al., 1998, Stiles and Jernigan, 2010, Urbán and Guillemot, 2014). Here, and building on our previous studies (Fu et al., 2017; Kim et al., 2022), we show instead that tuberal progenitors expand dramatically over the period HH10-HH13, resulting in the rapid separation of the *FOXG1*-expressing telencephalon and the *FOXA2/SHH* hypothalamic floor plate (Figure 1). Three lines of evidence indicate that tuberal progenitors are generated from anterior-most hypothalamic floor plate-like cells when BMP signalling is transiently activated in these cells.

First, at HH8-HH10, the appearance of pSMAD1/5/8 in anterior-most *FOXA2/SHH/NKX2-1*-expressing hypothalamic floor plate-like cells shortly precedes the upregulation of the tuberal progenitor markers, *SIX6/TBX2* (Figures 1, 2; Supplementary Fig 1). Thereafter, until HH17 (the latest stage examined), pSMAD1/5/8 is present immediately anterior to the anterior-most *FOXA2^(high)^/SHH*-expressing cells, which by this stage marks supramammillary hypothalamus and floor plate (Figure 2; Supplementary Figure 3). As the tuberal progenitor region lengthens, pSMAD1/5/8 always marks its posterior limit.

Second, fate-mapping studies show that tuberal hypothalamic cells transiently express the floor plate markers *FOXA2/SHH*, before upregulating tuberal progenitor markers (Figure 2; Supplementary Figure 2). A steady stream of anterior hypothalamic floor plate cells gives rise to tuberal progenitors, which migrate anteriorly relative to the *LHX3+* hypophysial placode/future Rathke’s pouch (Figure 2). Neighbouring cells always retain their relative positions along the anterior-posterior axis so that tuberal progenitors are laid down in an anterior-to-posterior order as they translocate anteriorly (Figure 2). For instance, progenitor cells that at HH10 express the posterior tuberal marker *TBX2* give rise to *SIX6*-expressing anterior tuberal progenitor cells at HH17, while *FOXA2*-positive floor plate-like cells at HH10 give rise to *TBX2*-positive posterior tuberal progenitor cells at HH17 (Figure 2; Supplementary Figure 2). Our studies build on previous fate-mapping studies of the HH10 embryo, which showed the anterior wards growth of *FGF10*-expressing progenitors that overlie the anterior prechordal mesoderm (Fu et al., 2017). Our new data suggests a progression from hypothalamic floor plate to posterior tuberal progenitor, anterior tuberal progenitor, anterior tuberal neurogenic progenitor, anterior neural precursor, and finally to mature tuberal neuron. Multi-label studies that reveal tuberal markers resolving into discrete spatial domains provide support for this sequence (Figure 2, Supplementary Figure 1) with *SIX6* becoming increasingly offset from its initial position with respect to *FOXA2*. While we cannot exclude that a small subset of anterior-most *SIX6*-expressing progenitors have a different history, our results indicate that most, if not all, tuberal *SIX6*-expressing anterior neurogenic progenitors transiently activate BMP Smads and express *TBX2*, and are derived from *FOXA2*-expressing hypothalamic floor plate-like cells.

Third, gain- and loss-of-function studies demonstrate that BMP signalling is necessary and sufficient for the induction of tuberal hypothalamic progenitors from floor plate-like cells. Our studies show that tuberal specification is initiated several hours before the appearance of tuberal markers, since isolated anterior-most hypothalamic floor plate cells from HH6 embryos switch floor plates for tuberal markers when cultured to a HH14-18 equivalent (Figure 4). Exposure of explants to the BMP inhibitor, Noggin, prevents this transition (Figure 5), and conversely, more posterior hypothalamic floor plate cells that normally fate-map to the posterior (mammillary) hypothalamus (Fu et al., 2017) can be induced to upregulate tuberal markers by ectopic BMP exposure (Figure 4). Notably, while early exposure to Noggin (at HH6) completely inhibits the tuberal progenitor programme, later exposure (at HH10) - both in *ex vivo* culture and *in vivo* - spares *SIX6*-expressing anterior tuberal progenitors, preventing only the development of posterior *TBX2*-expressing posterior tuberal progenitors (Figure 5).

A general question in development is how a limited number of extrinsic signals can robustly instruct neural progenitors to generate the immense diversity of cell types in the CNS. One intriguing possibility suggested by this work is that the conveyor belt-like movement of hypothalamic floor plate cells could provide a mechanism for temporal patterning of tuberal neurogenic progenitors, with the time at which hypothalamic floor plate-like cells are exposed to BMPs determining their future identity, as per the patterning of the body axes in zebrafish (Tucker et al., 2008). We detect changes in the gene expression profile of *FGF10*-expressing tuberal hypothalamic progenitors between HH10 and HH13, providing candidates for the molecular players behind the changing competence of these cells over time (Figure 7). The upregulation of *SOXE, ID*, and *ONECUT* family transcription factors, as well as differential expression of the *SHH, WNT, FGF* and *NOTCH* pathway components, could potentially determine the specification of distinct tuberal neuronal and glial subtypes as they encounter BMP signalling.

### A neuroepithelium-intrinsic BMP-SHH balance governs tissue homeostasis of tuberal progenitors

Many studies have shown the dependence of hypothalamic induction on the ventrally located prechordal mesoderm, which expresses BMP ligands (Dale et al., 1997, Dale et al., 1999, Mathieu et al., 2002, Ellis et al., 2015). Nonetheless, our *ex vivo* experiments indicate that, from as early as HH6, the tuberal differentiation programme can be sustained in the absence of any extrinsic sources of BMPs (Figure 4), suggesting the importance of a neuroepithelial-intrinsic mechanism.

Our *in vivo* experiments and previous studies (Corman et al., 2018, Fu et al., 2017, Manning et al., 2006, Trowe et al., 2013) suggest that a fine local balance between SHH and BMP signals sustains the tuberal progenitor programme. *TBX2* is a direct target of canonical BMP signalling (Shirai et al., 2009) and cell autonomously inhibits *SHH* expression in posterior tuberal progenitors (Figure 1) (Manning et al., 2006). Hence, *TBX2* downregulation and *SHH* upregulation may follow from the loss of BMP signalling as tuberal progenitors translocate anteriorly. Potentially, as in the mouse, a *SHH-SIX6* positive feedback loop (Jeong et al., 2008) may reinforce this molecular cascade, ensuring robust, BMP-dependent, anterior-posterior patterning (Figure 8, points 1-4).

**Figure 8:**
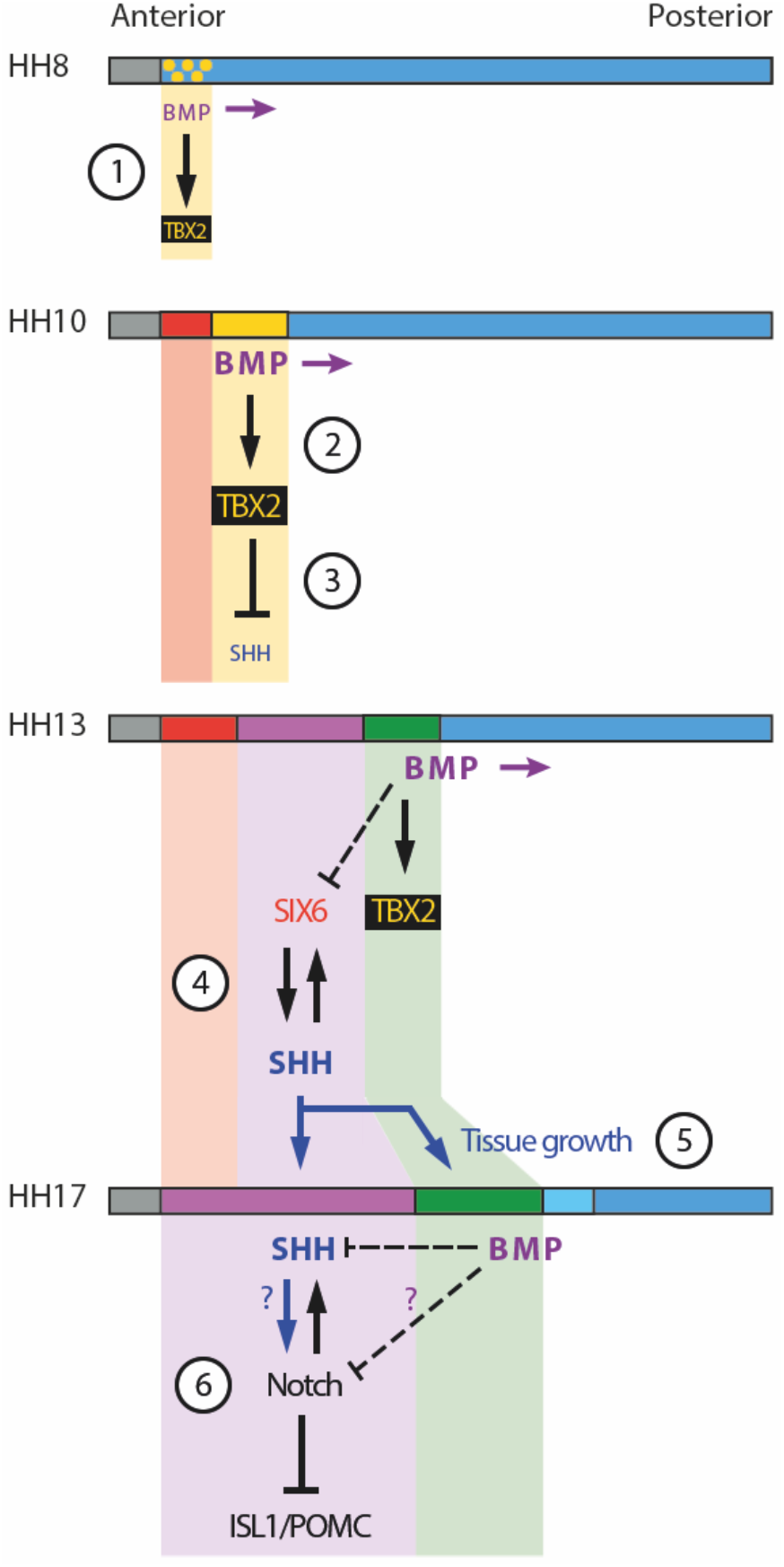
Schematic depicting selected signalling/transcription factor interactions in developing tuberal neuroectoderm. (1) The onset of BMP signalling in HH8 anterior neuroectoderm starts to induce *TBX2* in floor plate-like cells. (2) BMP signalling starts to move posteriorly within the neuroepithelium (purple arrows), promoting *TBX2* expression in its vicinity. (3) TBX2 in turn represses *SHH*, which starts to be downregulated. (4) A mutually reinforcing feedback loop between SIX6 and SHH may help to establish/maintain robust expression of both factors in the anterior tuberal domain. Note that BMP signalling represses *SIX6* expression after HH15, whether directly or indirectly (dotted line). (5) SHH produced by anterior tuberal progenitors may feed back onto posterior tuberal progenitors to sustain their growth, and that of the anterior domain. (6) By unknown mechanisms (question marks), finetuning of Notch signalling ensures the correct balance of proliferation versus neurogenesis in the tuberal hypothalamus. Horizontal coloured bars represent HH8-HH17 neuroectoderm; colouring is the same as Figure 1Q, Figure 2P. See Figure 2P for corresponding gene expression profiles.

Aside from their initial specification, our data shows that prolonged or excessive BMP levels disrupt later tuberal hypothalamic development, suggesting that BMP signalling must be switched off to progress to the anterior tuberal program and ensure the correct balance between growth and differentiation. BMP inhibition in HH15 explants led to the loss of posterior, and the gain of anterior tuberal markers (Figure 7). Prolonged exposure of nascent tuberal progenitors to BMPs at HH10 compromised anterior tuberal development at HH13 and led to a marginally larger *TBX2*-positive domain than in control embryos, consistent with BMPs inducing *TBX2* (Figure 6, (Shirai et al., 2009, Singh et al., 2009). At HH18, embryos show either a complete loss (Supplementary Figure 7) or a dramatic reduction (Figure 6) in *SIX6/SHH/NKX2-1-positive* anterior tuberal progenitors, as well as of *SHH*-dependent genes such as *NKX2-2* (Shimogori et al., 2010). In contrast to the effects seen at HH13, however, at HH18 posterior tuberal progenitor markers (*TBX2/FGF10/RAX/NKX2-1*) are also reduced or eliminated. Chick and mouse studies have shown that SHH signalling from anterior tuberal progenitors is required for the growth and differentiation of the neurogenic tuberal hypothalamus (Carreno et al., 2017, Corman et al., 2018, Fu et al., 2017, Shimogori et al., 2010, Szabó et al., 2009), and that a transient reduction in SHH signalling disrupts *FGF10* in posterior tuberal hypothalamic progenitors (Fu et al., 2017). We therefore propose that the small size of the posterior tuberal domain in BMP-treated embryos may be secondary to the loss of SHH deriving from anterior tuberal progenitors, and that, as in the developing cerebellum, SHH from more differentiated cells promotes proliferation of earlier-stage progenitors (Lewis et al., 2004, Wechsler-Reya and Scott, 1999) (Figure 8, point 5). Supporting this, the loss/reduction of tuberal markers seen following prolonged BMP exposure is accompanied by a significant decrease in proliferating cells (Figure 6).

Finally, despite the loss of tuberal progenitors in BMP-exposed embryos, neurogenic (*DLL1/ASCL1)* and tuberal neuronal markers (*ISL1/NR5A1/POMC*) were detected. This was furthermore associated with a reduction in *HES5*, suggesting that Notch signalling is also compromised in these embryos. This would be expected to result in premature neuronal differentiation of hypothalamic neurons (Aujla et al., 2013, Place et al., 2022, Ratié et al., 2013), providing a possible explanation for the presence of these markers even as progenitor development is heavily compromised. Our results overall suggest that an anterior-to-posterior wave of BMP signalling initiates first anterior, then later posterior, tuberal differentiation programs in floor plate-like neuroepithelial progenitors. The subsequent chain of molecular events establishes a precise spatiotemporal sequence of BMP, SHH, FGF and Notch signalling, ensuring the correct balance between progenitor specification, proliferation, and neurogenesis, laying the groundwork for the further development of this highly complex hypothalamic region (Figure 8, points 1-6).

### Establishment of a stem cell-like zone in the tuberal hypothalamus?

Our *ex vivo* studies show that tuberal patterning is largely complete by HH14, although anterior tuberal progenitor cells will continue to differentiate for many days thereafter. Our *in silico* bioinformatic analysis indicates that by HH20/21, posterior tuberal progenitor cells undergo a significant change, upregulating *VIM* and *FABP7*, markers of radial glial cells. Elsewhere in the CNS, the transition of neuroepithelial cells to radial glial cells coincides with the onset of stem-like character, and potentially, this is true also in the posterior tuberal hypothalamus. In all vertebrates examined to date, specialised radial glial-like cells, termed tanycytes, form the critical constituents of a stem cell-like niche that is present in the adult hypothalamus, situated in the median eminence, adjacent to the pituitary gland. Studies in the mouse have shown that adult *FGF10*-expressing tanycytes retain stem and progenitor-like activities (Goodman et al., 2020; Haan et al., 2013; Robins et al., 2013). We propose that the molecular changes that we detect in posterior tuberal progenitor cells over HH13-HH21 govern the transition from neurogenic *FGF10*-expressing neuroepithelial cells to *FGF10*-expressing radial glia, and the onset of stem-like characteristics in these cells. This work therefore indicates candidate regulators that govern development of the adult hypothalamic stem cell niche.

## Methods

### Chick Collection

Fertilized Bovan Brown eggs (Henry Stewart & Co., Norfolk, UK) were used for all experiments. All experiments were performed according to relevant regulatory standards (University of Sheffield). Eggs were incubated and staged according to the Hamburger-Hamilton chick staging system (Hamburger and Hamilton, 1992).

### Chicken HCR

Hamburger & Hamilton stage 8-20 embryos were harvested and fixed in 4% paraformaldehyde. HCR v3.0 was performed on embryos and cryosections using reagents and modified protocol from Molecular Instruments, Inc. Samples were preincubated with hybridization buffer for 30 min and the probe pairs were added and incubated at 37°C overnight. The next day samples were washed 4 times in the probe wash buffer, twice in the 5xSSC buffer, and preincubated in Amplification buffer for 5 min. Even and odd hairpins for each gene were snap-cooled by heating at 95°C for 90 sec and cooling to RT for 30 min. The hairpins were added to the samples in amplification buffer and incubated overnight at RT in the dark. Samples were then washed in 5xSSC and counterstained with DAPI. For multiplexing, after imaging with the first set of probes, wholemount samples were treated with DNAase (0.2 U/μl, Roche) overnight, washed 3 times in 30% formamide and 2xSSC and 3 times in 2xSSC. Slides were then preincubated with the hybridization buffer, the next set of probes were added, and the process was repeated.

### Immunohistochemistry

Embryos and explants were analysed by immunohistochemistry according to standard techniques (Manning et al., 2006). Embryos or cryosectioned sections were analysed using the following antibodies anti-pSMAD1/5/8 (1:500, Cell Signalling), anti-FOXA2 (1:50, DSHB), anti-Lhx3 (1:50, DSHB), anti-SHH (1:50, DSHB), anti-Islet1 (1:50, DSHB), anti-PH3 (1:1000, Cell Signalling). Secondary antibodies (1:500, Jackson Immunoresearch) were conjugated with anti-Alexa 488 or 594. Images were taken using a Zeiss Apotome.

### Single-cell analysis

We analysed our previous scRNA-Seq data for the developing chicken hypothalamus (Kim et al., 2022). Hypothalamic cells expressing *‘FGF10’* were subsetted and used in this study. Differential genes expressed in FGF10-positive cells across developmental time points were plotted on the heatmap.

### Explant culture

Explants of prospective hypothalamus were isolated from either HH6 or HH8 embryos by Dispase treatment and cultured in collagen beds (Ohyama et al., 2005). Explants were either treated with recombinant BMP2/7 heterodimers (32 nM, R&D Systems, Cat No. 3229-BM-010) or Noggin (300 ng/μl, R&D Systems, Cat No. 1967-NG) for 48 or 72 hrs, and processed for in situ hybridization chain reaction (HCR) or immunohistochemistry.

### *In vivo* manipulation of BMP signalling

BMP2/7 or Noggin soaked Affi-Gel beads were grafted into the anterior region of HH10 embryos. Embryos were allowed to develop to HH13 or HH17 (42 hrs), and processed for HCR and immunohistochemistry.

### Fate Mapping

HH10 embryos were windowed and a dorsal incision was made to access the ventral midline/floorplate. To aid visualisation, Coomassie Blue was diluted in L15 at 0.5 μl/ml and injected under the embryo using a 23G needle and syringe. 50 μg tubes of CellTracker™ CM-DiI Dye (Invitrogen, Cat No. C7000) were diluted in 30 μl ethanol, and 2 μl loaded into a fine glass needle, previously pulled to a sharp point using a needle puller. DiI was injected into embryos by hand using a Parker Picospritzer II set to 15 psi for 10-20 msec. Embryos were allowed to develop to HH17 and then harvested, fixed and processed for HCR and immunohistochemistry.

### Image acquisition and quantification

Fluorescent images were taken on a Zeiss Apotome 2 microscope with Axiovision software (Zeiss) or Leica MZ16F microscope or Olympus BX60 with Spot RT software v3.2 or Nikon W1 Spinning Disk Confocal with Nikon software. Images were acquired using a 4x (Leica), 10x (Leica and Zeiss) and 10x and 20xobjective Zeiss and Nikon microscope. Images were processed and digitally aligned using Image-J (FIJI) and Adobe Photoshop 2021. Unpaired *t*-tests were run on GraphPad Prism 9, and *p* < 0.05 was taken as significant. Mean ± SEM are plotted.

---

**Figure S1:**
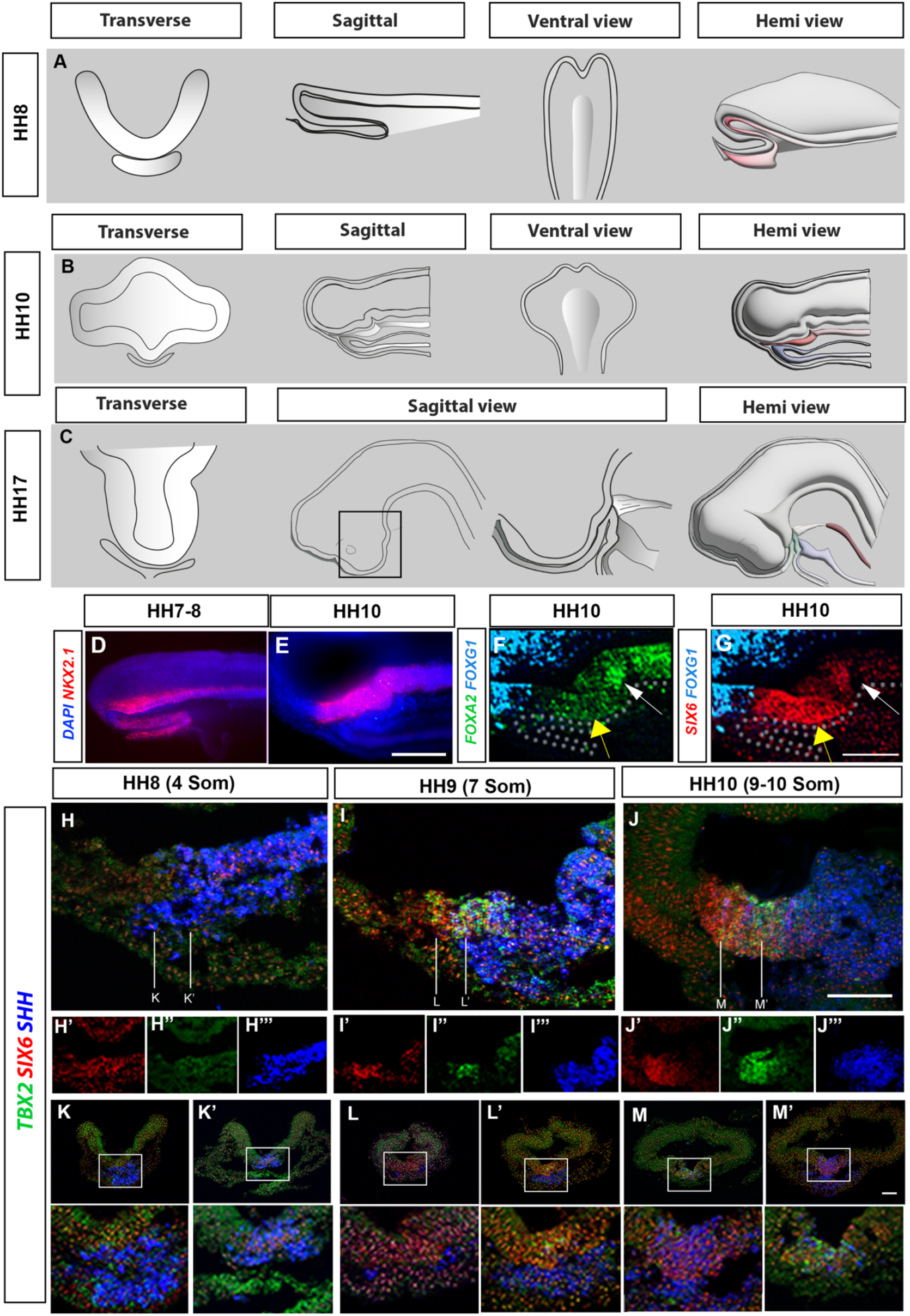
Profiling developing tuberal progenitors. (A-C) Schematics depict transverse sections, sagittal sections, ventral wholemount and hemi-dissected views of HH8 (A), HH10 (B) and HH17 (C) embryos. (D-E) Maximum intensity projections of hemi-views at HH8-HH10 after HCR for *NKX2-1*. (F, G) Maximum intensity projections of hemi-views at HH10 after HCR for *FOXG1, FOXA2, SIX6*. Same embryo as shown in Figure 1B, but *FOXA2* and *SIX6* channels separated. Yellow arrows show *SIX6/FOXA2^(low)^* region, white arrows show *FOXA2^(high)^* region. (H-J’’’) Maximum intensity projections of sagittal sections at HH8-HH10 after HCR for *TBX2, SIX6, SHH*. Embryos in H-J are same as in Figure 1L-M, but channels are separated in H’-J”’. (K-M’) Transverse views of anterior-most tuberal regions (approximate levels shown in H-J), after HCR for *TBX2, SHH and SIX6* at HH8, HH9 and HH10, and enlarged views of the boxed areas. All scale bars = 100μm

**Figure S2:**
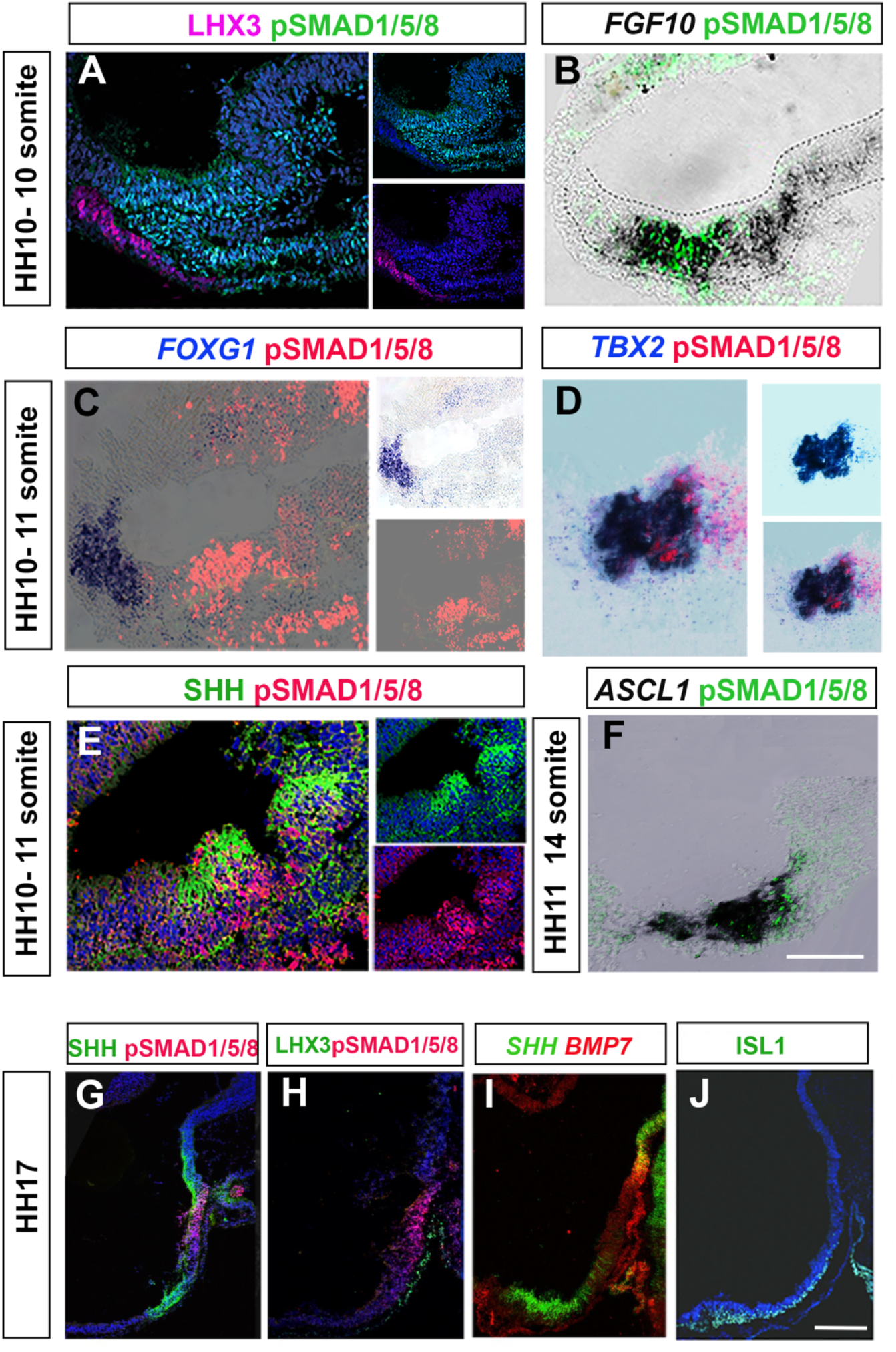
pSmad1/5/8-positive cells mark the posterior-most tuberal domain. (A) Sagittal section of a HH10 embryo immunolabelled with LHX3 and pSMAD1/5/8. pSmad1/5/8-expressing cells occupy the flat anterior ventral neuroepithelium. (B-D) Sagittal sections of HH10 embryos analysed by chromogenic i*n situ* hybridisation for *FGF10 (B), FOXG1 (C)* or *TBX2 (D)* and then immunolabelled to detect pSMAD1/5/8. pSmad1/5/8 shows extensive overlap with *FGF10*, does not overlap with *FOXG1*, and overlaps/lies just posterior to *TBX2*. (E) Sagittal section of HH10 embryo immunolabelled for SHH and pSMAD1/5/8. The two markers show a similar anterior boundary. (F) Sagittal section of a HH11 embryo analysed by chromogenic *in situ* hybridization for *ASCL1* and then immunolabelled to detect pSMAD1/5/8. pSMAD-positive nuclei are posterior to *ASCL1*-expressing regions. (G-J) Serial adjacent sagittal sections taken through a HH17 embryo, immunolabelled to detect SHH and pSMAD1/5/8 (G), LHX3 and pSMAD1/5/8 (H), ISL1 (J) or analyzed by HCR to detect *SHH* and *BMP7* (I) . All scale bars = 100μm. Each panel shows a representative image from a minimum n= 5 embryos/ label combination

**Figure S3:**
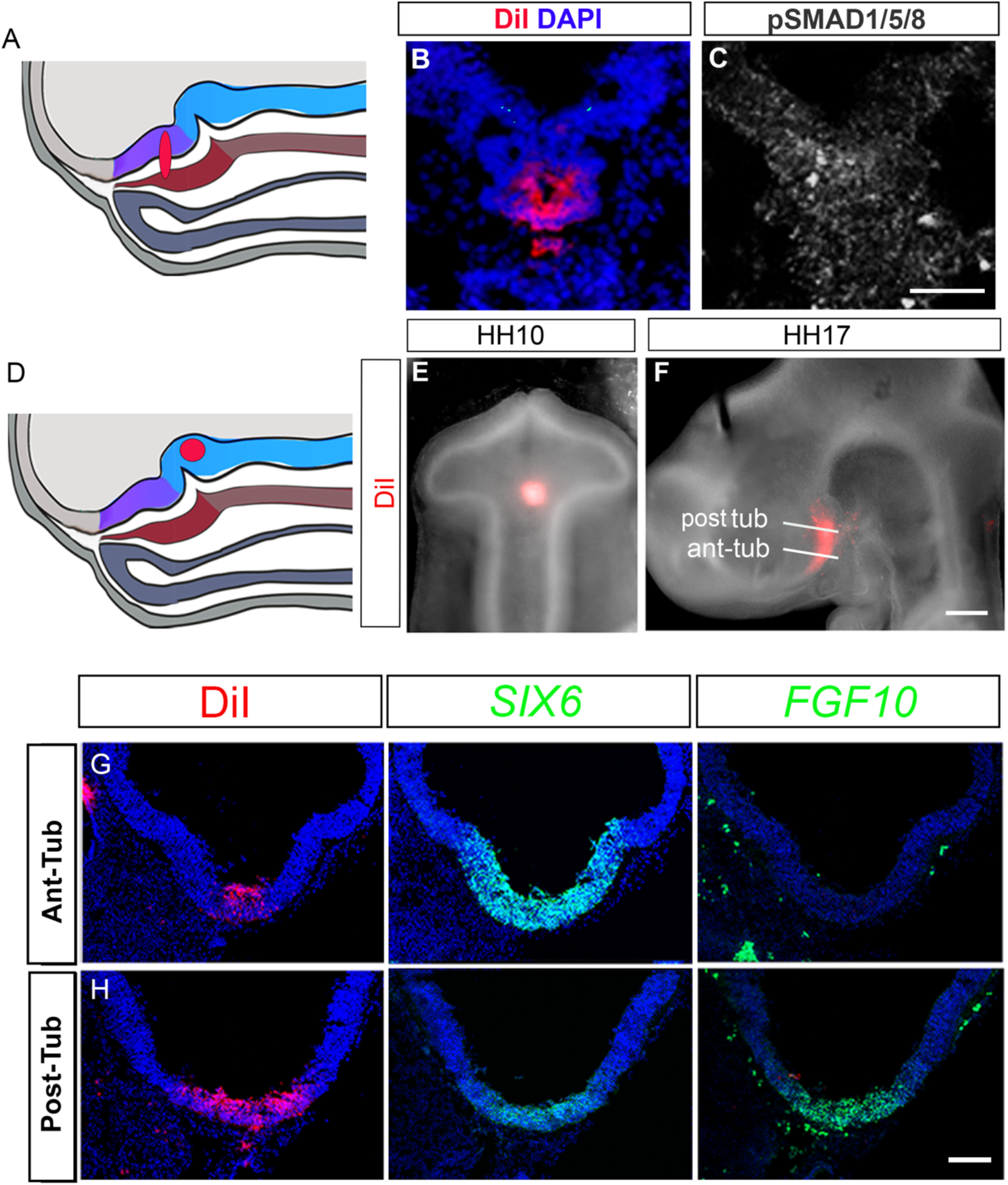
Fate-mapping shows the origin of tuberal hypothalamic cells. (A) Schematic depicts a DiI injection at HH10 that labels both the pSmad1/5/8-positive neuroepithelium (purple), and underlying prechordal mesoderm (dark brown). (B, C) Serial adjacent sections of a HH10 embryo targeted as in (A). DiI is restricted to the flat neuroepithelium, at the base of a characteristic fold (B), within the pSmad1/5/8-positive domain (C). (D, E) Schematic depicts embryo in which DiI was injected in the region occupied by hypothalamic floor plate cells. (F) Wholemount view of embryo in (E) after 48 hours incubation to HH17. DiI is mainly localised to the posterior tuberal hypothalamus (post-tub), petering out in the anterior tuberal (ant-tub) hypothalamus. (G-H) Transverse sections through the same embryo as in (F); plane of sections marked by white lines, analysed by HCR to detect *SIX6* and *FGF10*. There is extensive DiI labelling in posterior tuberal progenitors (FGF10/SIX6^(low)^, petering out in anterior tuberal progenitors (*SIX6^(high)^, FGF10*-negative). All scale bars = 100μm.

**Figure S4:**
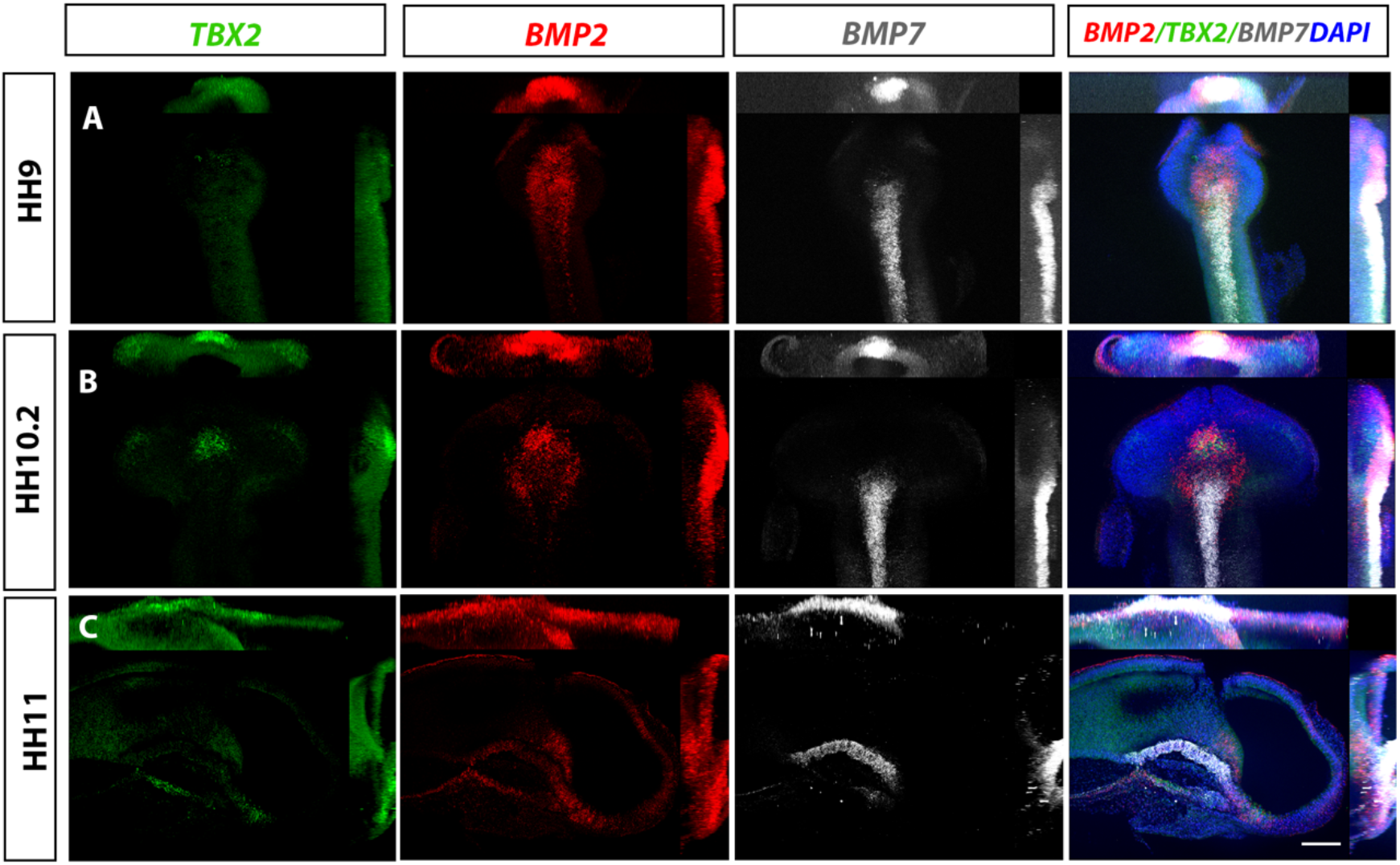
Tuberal progenitors arise in and around BMP-expressing neuroepithelial cells. (A-C) Maximum intensity projections of ventral (A, B) and hemi-views (C) of HH9-HH11 embryos, analysed by HCR for *TBX2, BMP2*, and *BMP7*. Individual channels are shown in left hand panels, and combined views are shown on the right hand panels. (A) At HH9, *BMP2* is expressed as a cap anterior to/around *BMP7*-expressing cells. In this embryo, *TBX2* expression is not detected. (B, C) At HH10-HH11, *TBX2* is upregulated in neuroectoderm cells that express high levels of *BMP2*. All scale bars = 100 μm. All panels show representative images from n = 5 embryos

**Figure S5:**
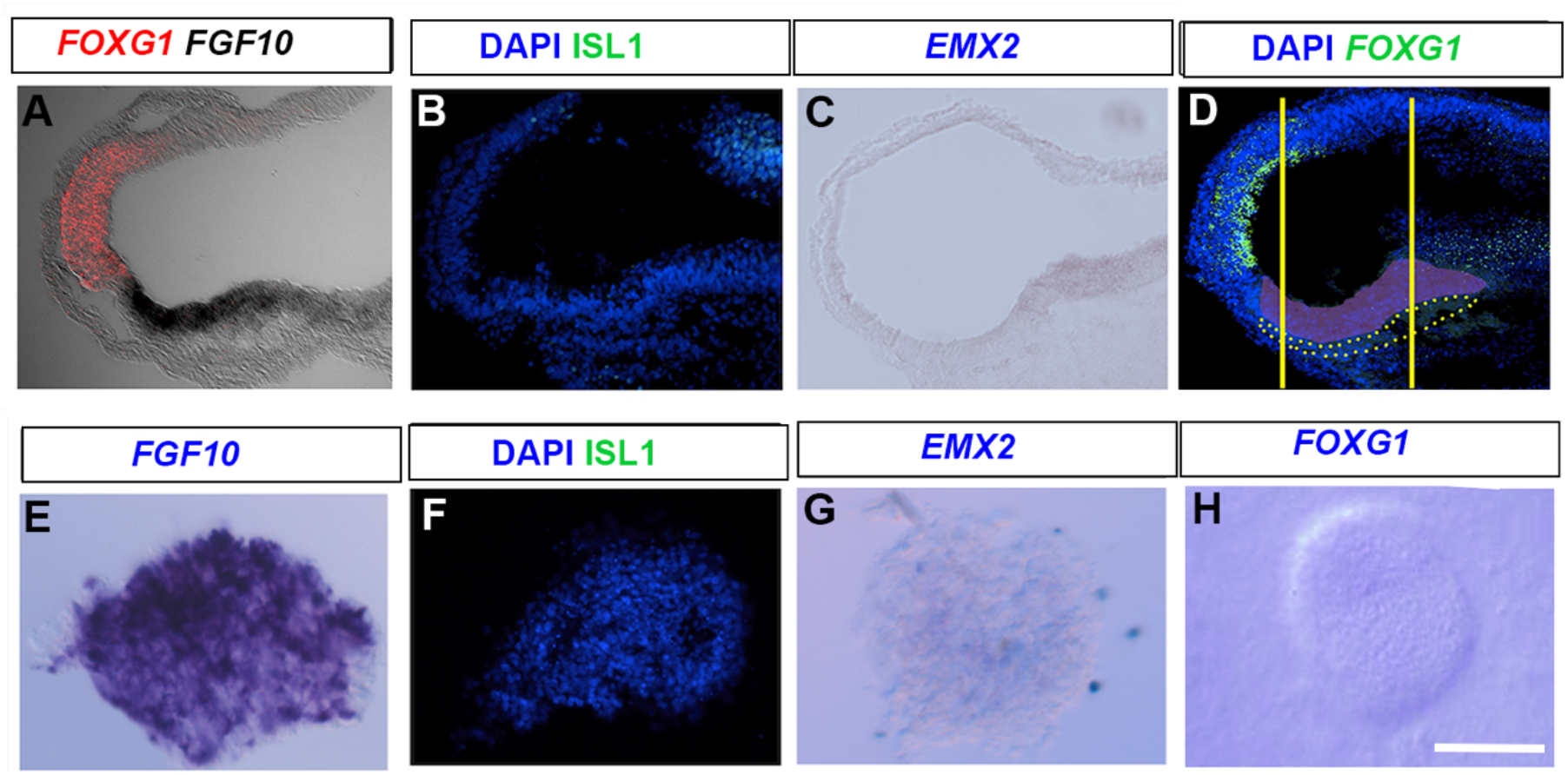
Accurate isolation of hypothalamic tissue. (A-D) Sagittal sections of HH10 embryos, analysed by chromogenic *in situ* hybridisation or immunolabelling to detect *FOXG1/FGF10*, ISL1, and *EMX2*. Pink region in D depicts emerging tuberal hypothalamus, and yellow lines show the region dissected for explant culture. (E-H) Sections through HH10 hypothalamic explants, isolated as in Figure 3, and examined at t=0hrs by chromogenic *in situ* hybridisation or immunolabelling. Explants express *FGF10* but not *FOXG1*, ISL1, or *EMX2*. All scale bars = 100 μm. All panels show representative images from a minimum of 6 embryos and 4 explants.

**Figure S6:**
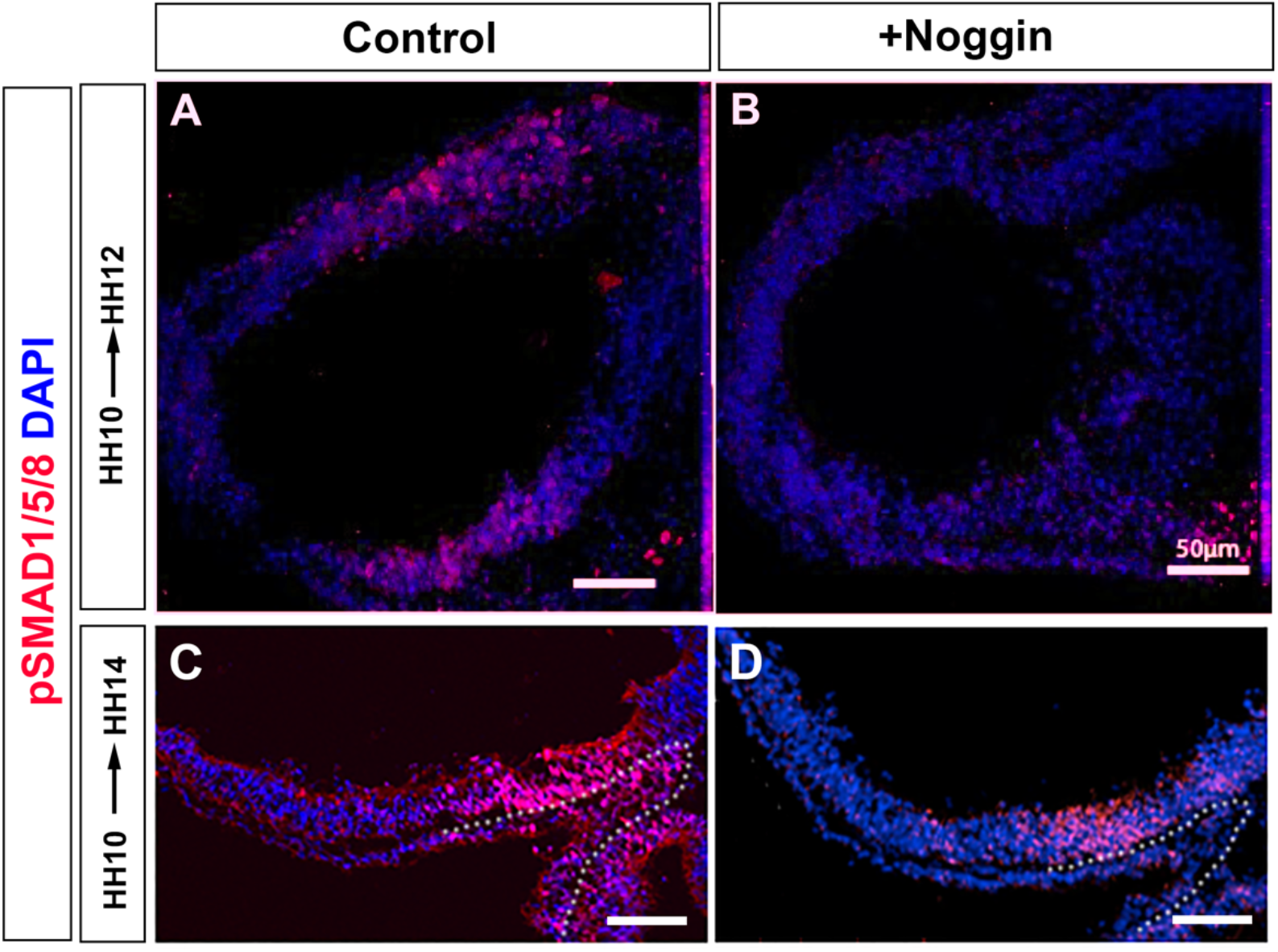
Transient reduction of pSMAD1/5/8 *in vivo*. (A-D) Sagittal sections taken through HH12 (A, B) or HH14 (C, D) embryos after implantation of a PBS-soaked control bead (A, C) or a Noggin-soaked bead at HH10, analysed by immunolabelling with pSMAD1/5/8 (B, D). All scale bars = 50μm Panels show representative images from n=3 embryos/time point/condition.

**Figure S7:**
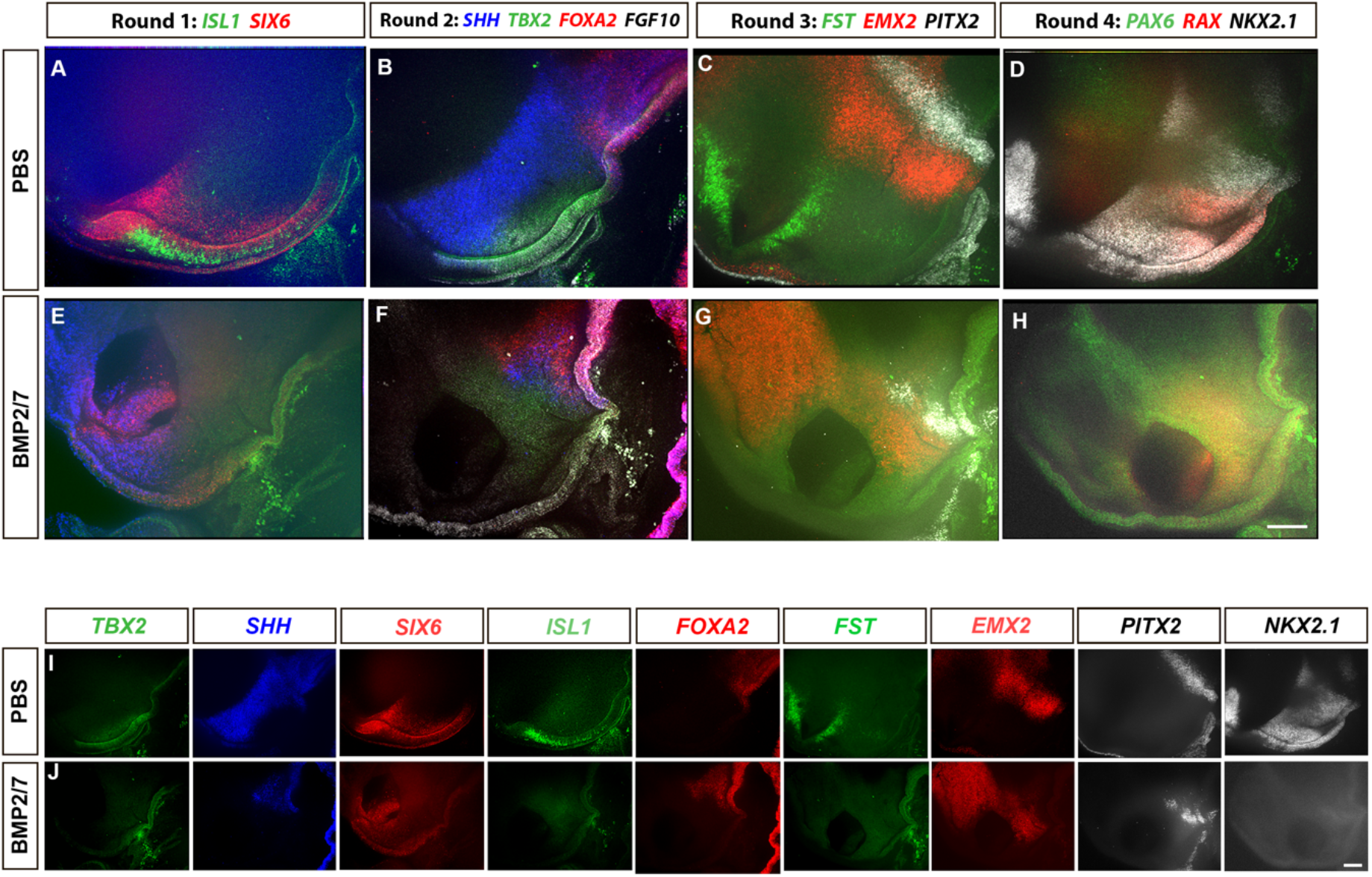
Ectopic BMPs eliminate tuberal cells. (A-D) Hemi-view of PBS bead-grafted control embryo after 4 rounds of multiplex wholemount HCR for SIX6, *ISL1* (Round1, A); *SHH*, *TBX2, FGF10*, and *FOXA2* (Round 2, B); *FST*, *PITX2*, and *EMX2* (Round 3, C) and *PAX6, RAX*, and *NKX2.1* (Round 4, D). (E-H). Hemi-view of BMP2/7-soaked bead grafted embryo after 4 rounds of multiplex wholemount HCR for *SIX6, ISL1* (Round1, E); *SHH, TBX2, FGF10*, and *FOXA2* (Round 2, F); *FST*, *PITX2*, and *EMX2* (Round 3, G) and *PAX6, RAX*, and *NKX2.1* (Round 4, H). (I-J) Shows individual channel view of the same control and BMP2/7-treated embryo shown above. n=2 embryos/condition. All scale bars = 100μm.

## ACKNOWLEDGMENTS

We thank Transcriptomics and Deep Sequencing Core (Johns Hopkins) for sequencing of scRNA-seq libraries. This work was supported by the Wellcome Trust (212247/Z/18/Z) to M.P.

